# Shotgun scanning glycomutagenesis: a simple and efficient strategy for constructing and characterizing neoglycoproteins

**DOI:** 10.1101/2020.06.28.176198

**Authors:** Mingji Li, Xiaolu Zheng, Sudhanshu Shanker, Thapakorn Jaroentomeechai, Tyler D. Moeller, Sophia W. Hulbert, Ilkay Koçer, Josef Byrne, Emily C. Cox, Qin Fu, Sheng Zhang, Jason W. Labonte, Jeffrey J. Gray, Matthew P. DeLisa

## Abstract

As a common protein modification, asparagine-linked (*N-*linked) glycosylation has the capacity to greatly influence the biological and biophysical properties of proteins. However, the routine use of glycosylation at naïve sites as a strategy for engineering proteins with advantageous properties is currently limited by our inability to construct large collections of glycoproteins for interrogating the structural and functional consequences of glycan installation. To address this challenge, we describe a combinatorial strategy termed shotgun scanning glycomutagenesis (SSGM) in which DNA libraries encoding all possible glycosylation site variants of a given protein are constructed and subsequently expressed in glycosylation-competent bacteria, thereby enabling rapid determination of glycosylatable sites in the protein. Moreover, the resulting neoglycoproteins can be readily subjected to available medium- to high-throughput assays, making it possible to systematically investigate the structural and functional consequences of glycan conjugation along a protein backbone. The utility of this approach was demonstrated with three different acceptor proteins, namely bacterial immunity protein Im7, bovine pancreatic ribonuclease A, and a human anti-HER2 single-chain Fv antibody, all of which were found to tolerate *N-*glycan attachment at a large number of positions and with relatively high efficiency. The stability and activity of many glycovariants was measurably altered by the *N-*linked glycan in a manner that critically depended on the precise location of the modification. Importantly, we anticipate that our workflow for creating and characterizing large ensembles of neoglycoproteins should provide access to unexplored regions of glycoprotein structural space and to custom-made glycoproteins with desirable properties.

## Introduction

Glycosylation of asparagine residues is one of the most abundant and structurally complex protein post-translational modifications (1, 2) and occurs in all domains of life (3). Owing to their relatively large size and hydrophilicity or simply their presence at definite locations, asparagine-linked (*N*-linked) glycans can significantly alter protein properties including biological activity, chemical solubility, folding and stability, immunogenicity, and serum half-life (4, 5). Hence, glycosylation effectively increases the diversity of the proteome by enriching the repertoire of protein characteristics beyond that dictated by the twenty canonical amino acids. For example, accumulating evidence indicates that the immune system diversifies the repertoire of antigen specificities by exclusively targeting the antigen-binding sites of immunoglobulins (IgGs) with post-translational modifications, in particular *N*-linked glycosylation (6). Moreover, the profound effect of glycans on proteins has prompted widespread glycoengineering efforts to rationally manipulate key glycosylation parameters (*e.g.*, glycan size and structural composition, glycosite location and occupancy) as a means to optimize protein traits for a range of different industrial and therapeutic applications (7–10).

Despite some notable successes, the routine use of glycosylation as a strategy for engineering proteins with advantageous properties is currently limited by our inability to predict which sites within a protein are glycosylatable and how glycosylation at permissive sites will affect protein structure and function. Indeed, a deeper understanding of the design rules (*i.e.*, how glycans influence the biological and biophysical properties of a protein) represents a grand challenge for the glycoprotein engineering field. To this end, computational approaches have enabled *in silico* exploration of glycosylation-induced effects on protein folding and stability (11, 12); however, these involve a trade-off between molecular detail and glycoprotein size, with full-atomistic molecular dynamics simulations typically limited to only short glycopeptides or protein domains (11). To experimentally probe the consequences of glycosylation ideally requires access to large collections of chemically defined glycoproteins in sufficient quantities for characterization (13). Mammalian cells represent an obvious choice to source proteins with both natural and naïve glycosites; however, studies using mammalian cell-based expression systems typically involve only a small number of designs (~15 or fewer) (14–17) presumably because of the time-consuming, low-throughput nature of gene transfection and culturing of mammalian cells. In addition, the intrinsic variability with respect to the glycan structure at a given site (microheterogeneity) can be unpredictable and difficult to control in mammalian expression systems. Another option is chemical synthesis, which can furnish structurally uniform glycopeptides for investigating the local effects of *N*-linked glycans on peptide conformation (18). While this approach is not amenable to full-length proteins, advances in expressed protein ligation (EPL) have opened the door to convergent assembly of chemically synthesized glycopeptides with recombinantly expressed protein domains to form larger glycoproteins bearing complex *N*-glycans installed at discrete sites (19). Using this technology, Imperiali and colleagues created a panel of seven site-specifically glycosylated variants of the bacterial immunity protein Im7 modified with the disaccharide *N*,*N*’-diacetylchitobiose (GlcNAc_2_) and assessed the kinetic and thermodynamic consequences of glycan installation at defined locations (20). Unfortunately, EPL is a technically demanding procedure, requiring manual construction of each individual glycoprotein, which effectively limits the number of testable glycosite designs to just a small handful.

To move beyond these “one-glycosite-at-a-time” methods for supplying glycoproteins, herein we describe a scalable technique called <u>s</u>hotgun <u>s</u>canning <u>g</u>lyco<u>m</u>utagenesis (SSGM) that involves design and construction of combinatorial acceptor protein libraries in which: (i) each member of the library carries a single *N-* glycosite “mutation” introduced at a defined position along the protein backbone; and (ii) the complete ensemble of glycan acceptor sites (sequons) in the library effectively covers every possible position in the target protein (**Fig. 1**). The resulting SSGM libraries are expressed using *N-*glycosylation-competent bacteria in the context of glycoSNAP (<u>glyco</u>sylation of <u>s</u>ecreted *N*-linked <u>a</u>cceptor <u>p</u>roteins), a versatile high-throughput screen based on extracellular secretion of glycosylated proteins (21). Using this new glycoprotein engineering tool, we constructed and screened SSGM libraries corresponding to three model proteins: bacterial immunity protein Im7, bovine pancreatic ribonuclease A (RNase A), and a human single-chain variable fragment antibody specific for HER2 (scFv-HER2). Our results revealed that installation of *N-*glycans was tolerated at a large number of positions and in all types of secondary structure, with relatively high *N-*glycosylation efficiency in the majority of cases. For many of these glycoproteins, the presence of *N-* glycans at naïve sites had a measurable effect on protein stability and/or activity in a manner that depended on the precise location of the modification. Taken together, these findings demonstrate the ability of the SSGM method to yield large collections of discretely modified neoglycoproteins that collectively reveal glycosylatable sites and provide insight on the influence that site-specific *N-*glycan installation has on structural and/or functional properties.

**Figure 1.**
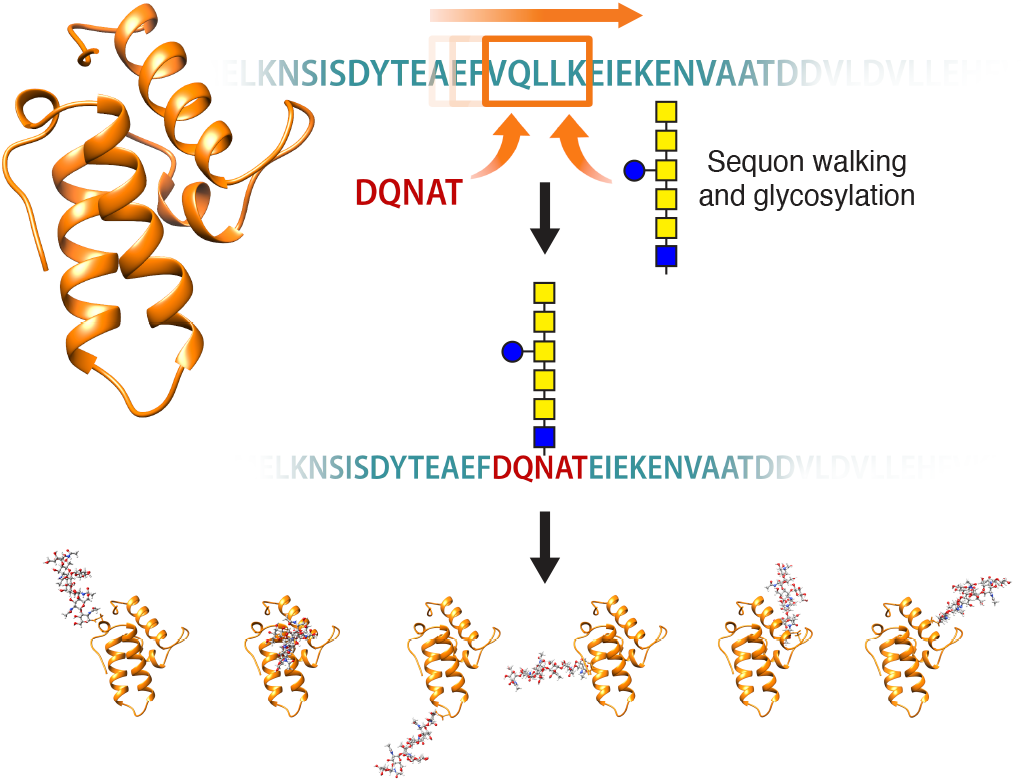
Constructing neoglycoproteins by shotgun scanning glycomutagenesis (SSGM). Schematic of SSGM, a glycoprotein engineering method based on combinatorial protein libraries in which glycosylation “sequon walking” is used to introduce an acceptor site at every possible position along a protein backbone. Note that the multi-residue nature of a sequon (*e.g.*, N-X-S/T or D/E-X_1_-N-X_2_-S/T where X, X_1_, X_2_ ≠ P) necessitates insertion or replacement of up to five additional amino acid substitutions at each position. The resulting library is expressed in glycoengineered bacteria, providing an opportunity for each library member to be expressed and glycosylated in a manner that is compatible with high-throughput screening via glycoSNAP to interrogate the glycosylation phenotype of individual variants. By integrating expressed SSGM libraries with multiplexable assays, the biochemical and biophysical properties of each neoglycoprotein can be individually interrogated.

## Results

### Reliable detection of acceptor protein glycosylation by glycoSNAP screening

To enable screening of SSGM libraries, we first sought to adapt glycoSNAP screening for proteins of interest (POIs). In the original glycoSNAP assay, we genetically modified *Escherichia coli* YebF, a small (10 kDa in its mature form) extracellularly secreted protein (22), with an artificial glycosite (*e.g.*, N-X-S/T or D/E-X_1_-N-X_2_-S/T where X, X_1_, X_2_ ≠ P) at its C-terminus. The modified YebF protein was expressed in *E. coli* cells carrying the *Campylobacter jejuni N-*glycosylation machinery (23) that were bound to a nitrocellulose filter membrane. Following secretion out of filter-bound colonies, putatively glycosylated YebF was captured on a second nitrocellulose membrane, which was probed with antibodies or lectins to detect *N-*linked glycans. In this way, glycoSNAP creates a convenient genotype–glycophenotype linkage for facile scoring (glycosylated versus aglycosylated) of YebF proteins secreted from individual bacterial colonies (**Fig. 2a**). Here, we hypothesized that genetic fusion of glycosite-modified POIs to YebF would result in extracellular secretion of the fusion protein such that glycans installed on the POI could be detected by the nitrocellulose membrane-based screening strategy. To test this hypothesis, we initially focused on *E. coli* Im7 as the POI for several reasons: (i) it is a small, globular 87-residue protein that lacks disulfide bonds and is well expressed in the periplasm where bacterial *N-*glycosylation occurs (23); (ii) although not a native glycoprotein, Im7 modified at its C-terminus with a DQNAT glycosylation tag can be glycosylated by the *C. jejuni N-*glycosylation machinery in *E. coli* (23); (iii) crystal structures are available for wild-type (wt) Im7 (24) and for Im7 in complex with its cognate toxin colicin E7 (ColE7) (25); and (iv) a limited set of seven Im7 variants was previously generated to determine the effects of GlcNAc_2_ attachment on folding and stability (20), providing some useful reference points for comparison.

**Figure 2.**
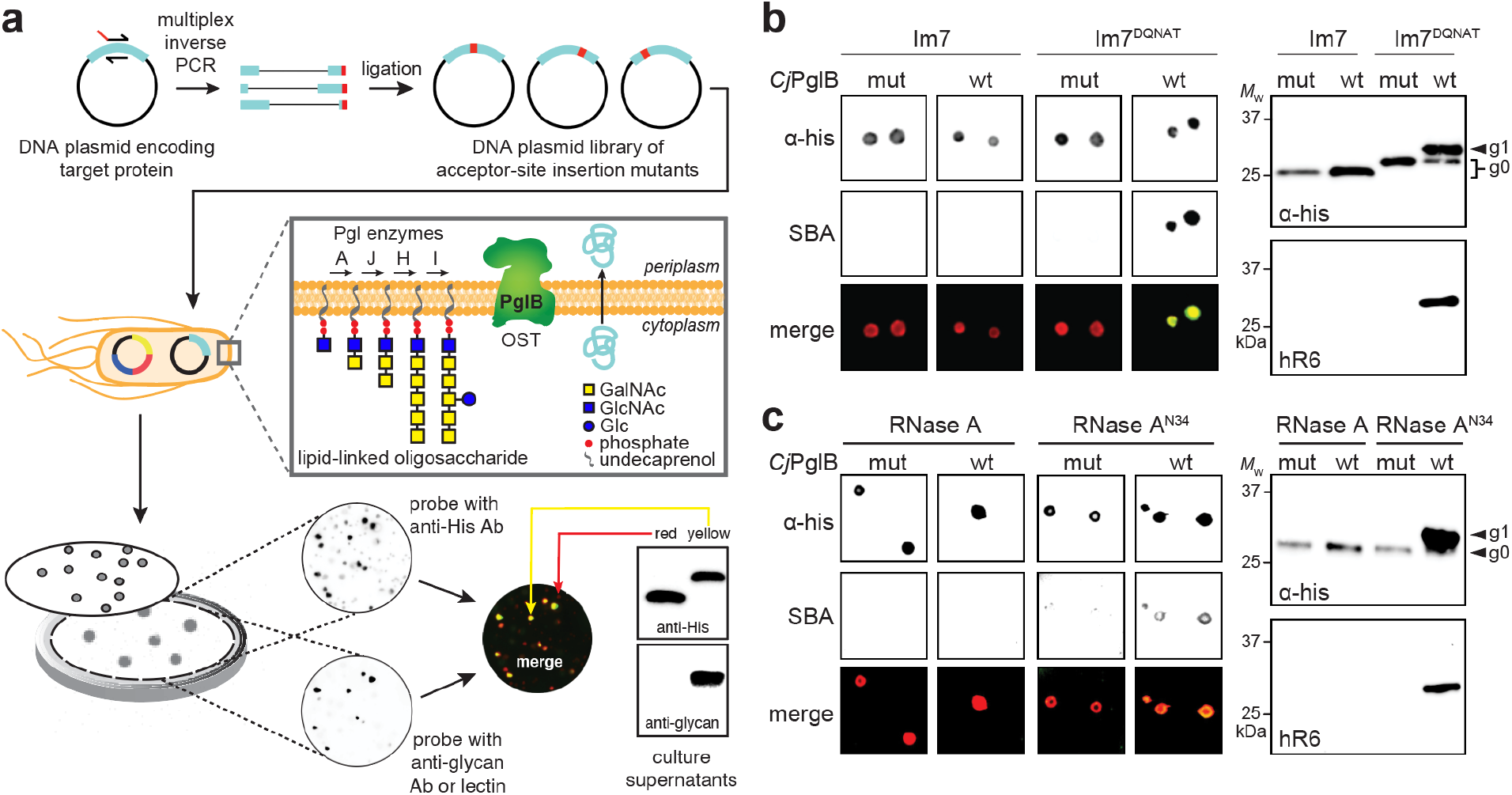
Construction and interrogation of SSGM libraries. (a) Schematic of SSGM library construction using multiplex inverse PCR. The resulting DNA plasmid library, encoding neoglycoprotein variants with glycosite substitutions at every possible position, was used to co-transform *E. coli* strain CLM24 along with two additional plasmids encoding the requisite *N-*glycosylation machinery from *C. jejuni*. The resulting bacterial library was plated on solid agar, after which colonies and their secreted glycoproteins were replica plated on nitrocellulose membranes as described in the text. (b) Immunoblot analysis of acceptor proteins in colony secretions (left) and extracellular supernatant fractions (right) derived from *E. coli* CLM24 carrying a plasmid encoding either YebF-Im7 or YebF-Im7^DQNAT^ along with plasmids encoding *N-*glycosylation machinery with either wild-type *Cj*PglB (wt) or an inactive mutant (mut). (c) Same as in (b) but with YebF-RNase A and YebF-RNase A^N34^ in colony secretions (left) and periplasmic fractions (right). Blots were probed with anti-polyhistidine antibody (α-His) to detect acceptor proteins and SBA or hR6 serum to detect glycans. Bottom color panels in (b) and (c) depict overlay of α-His and SBA blots (merge). Arrows denote aglycosylated (g0) and singly glycosylated (g1) forms of YebF-Im7^DQNAT^ or YebF-RNase A^N34^. Molecular weight (*M*_W_) markers are indicated at left. Results are representative of at least three biological replicates.

To determine whether Im7 was compatible with the glycoSNAP procedure, *E. coli* strain CLM24 was transformed with a plasmid encoding YebF-Im7 that was modified with a DQNAT glycosylation tag (26) at the C-terminus of Im7 along with two additional plasmids, one encoding glycosyltransferase (GT) enzymes for the biosynthesis of the *N*-glycan and the other encoding the oligosaccharyltransferase (OST) for transfer of the resulting *N-*glycan to acceptor proteins. To minimize microheterogeneity so that modified acceptor proteins all carried identical glycans, we created a system for producing homogeneous *N*-glycans with the structure GalNAc_5_(Glc)GlcNAc, which is one of several structurally related glycan donors that can be efficiently transferred to target proteins in *E. coli* by the *C. jejuni* OST PglB (*Cj*PglB) (27, 28). While the biotechnological value of this glycan is questionable, it served as an excellent model for our proof-of-concept SSGM studies for several reasons. First, it involves formation of the key GlcNAc-Asn linkage, which is the same as found in prototypic eukaryotic *N*-glycans. Second, it has the potential to be remodeled as a complex-type eukaryotic glycan via a two-step enzymatic trimming/transglycosylation process (29). Third, its structural uniformity and relative abundance when produced heterologously in *E. coli* cells, as well as its compatibility with PglB, all help to ensure that differences in glycosylation efficiency are minimally affected by substrate-related factors and are instead attributable to accessibility of a given acceptor site.

When plated on solid agar and subjected to the colony-blotting method, cells expressing YebF-Im7_DQNAT_, or a control YebF-Im7 construct that lacked the glycosylation tag, were able to secrete the fusion into the extracellular medium as evidenced by cross-reaction of an anti-His antibody with the membranes (**Fig. 2b**). However, only the strain expressing YebF-Im7^DQNAT^ in the presence of wt *Cj*PglB, but not a *Cj*PglB variant rendered inactive by two active-site mutations (D54N and E316Q) (21), gave rise to colonies that reacted with soybean aggluntinin (SBA) (**Fig. 2b**), a lectin that binds terminal GalNAc residues in the *C. jejuni N-*glycan (27). The colony blotting results were corroborated by immunoblot analysis of culture supernatants, which revealed that YebF-Im7 and YebF-Im7^DQNAT^ were both secreted into the extracellular medium but only the latter was glycosylated as evidenced by the appearance of a higher molecular weight band in the blot probed with glycan-specific antiserum (**Fig. 2b**). As expected, no glycan-specific signal was detected in colony blots or immunoblots corresponding to cells carrying the mutant *Cj*PglB enzyme (**Fig. 2b**). Importantly, the predominant glycan attached to YebF-Im7^DQNAT^ corresponded to GalNAc_5_(Glc)GlcNAc, which represented >98% of all detected glycoforms as confirmed by mass spectrometry (**Supplementary Fig. 1**). Collectively, these results confirmed the compatibility of bacterial Im7 with our glycosylation workflow, yielding homogenously modified acceptor proteins that were readily detected by glycoSNAP screening.

### Rapid identification of acceptor site permissiveness using SSGM

Next, the plasmid encoding YebF-Im7 was mutagenized to create a library of Im7 gene sequences, each carrying an individual sequon substitution and cumulatively covering all positions in the Im7 protein. Mutagenesis was performed using multiplex inverse PCR (30) with a set of divergent abutting primers that were designed to amplify the entire plasmid and introduce an acceptor asparagine residue at every position in the Im7 gene (with the two upstream and two downstream residues being changed to DQ and AT, respectively), thereby yielding a highly focused plasmid library enriched with in-frame clones each bearing a single DQNAT acceptor motif at a defined position (**Fig. 2a**). Indeed, next-generation sequencing of the pre-selected plasmid library confirmed complete sequence coverage for all glycosite positions in Im7, with >10^3^ reads detected for all but one position. (**Supplementary Fig. 2**). With all glycosite variants present and accounted for, the resulting plasmid library was introduced into strain CLM24 carrying the requisite *N-* glycosylation machinery and the library-transformed cells were plated on solid agar and subjected to glycoSNAP screening. From one membrane, we detected a total of ~200 glycosylation-positive colonies, of which 20 were randomly chosen for further analysis. Sequencing confirmed that a single in-frame DQNAT motif was present in each isolated hit, with the Im7^N37^ and Im7^N58^ variants (where the superscript denotes the location of the asparagine residue) occurring three and two times, respectively (**Fig. 3a**). The hits were fairly evenly distributed throughout the entire Im7 sequence and situated in every type of secondary structure including bends, turns, and α-helices, consistent with X-ray crystallographic data showing that occupied glycosylation sites can occur on all secondary structural elements (31). Immunoblot analysis confirmed that each of the selected clones was efficiently glycosylated (**Fig. 3b**).

**Figure 3.**
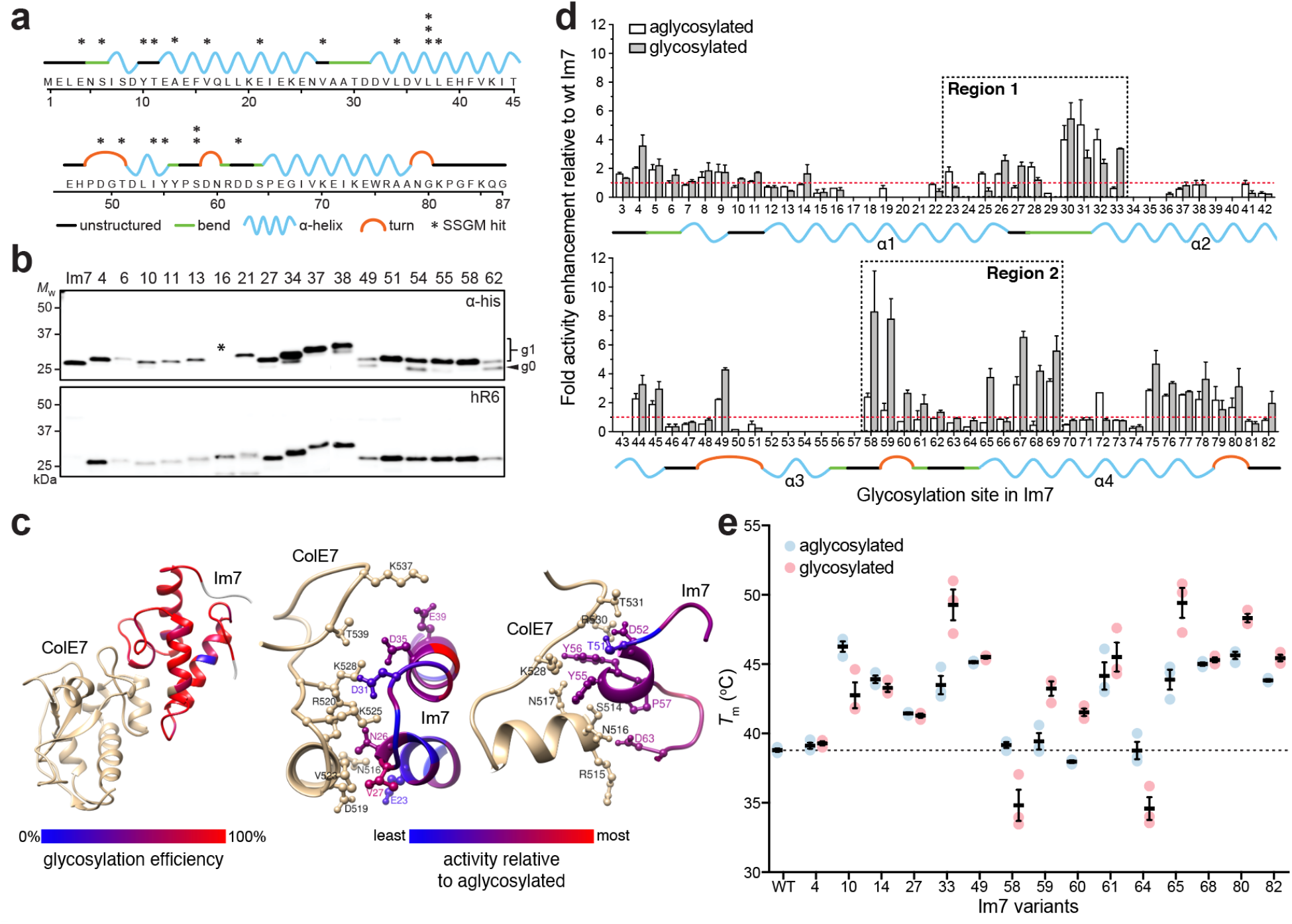
Construction and characterization of bacterial Im7 neoglycoprotein library. (a) Primary sequence and predicted secondary structure for *E. coli* Im7 immunity protein. Asterisks denote location and frequency of glycosite hits isolated using SSGM. Predicted structures adapted from PDB ID 1AYI. (b) Immunoblot analysis of supernatant fractions from CLM24 cells carrying plasmids encoding YebF-Im7 fusions with sequon mutations at indicated position and requisite *N-*glycosylation machinery. Blots were probed with anti-polyhistidine antibody (α-His) to detect acceptor protein (top panel) and hR6 serum against the glycan (bottom panel). Markers for aglycosylated (g0) and singly glycosylated (g1) forms of acceptor proteins are indicated at right. Molecular weight (*M*_W_) markers are indicated at left. Asterisk indicates construct with mutation that introduced stop codon just before 6xHis tag, preventing α-His detection. Results are representative of at least three biological replicates. (c) Mapping of cell-based glycosylation efficiency onto three-dimensional structure of Im7 in complex with ColE7 (left). Heatmap analysis of the glycosylation efficiency was determined based on densitometric quantification of the percent glycosylated (defined as g1/[g0+g1] ratio) for each acceptor protein in the anti-His immunoblot. Detailed interactions between ColE7 and Im7, highlighting sidechains of Im7 in the regions of α1-loop12-α2 (residues 19-39; middle) and loop23-α3-loop34 (residues 46-63; right). Heatmap analysis of change in binding activity was determined by normalizing activity measured for glycosylated sequon variant by aglycosylated counterpart. (d) Binding activity of glycosylated (gray bars) and aglycosylated (white bars) YebF-Im7 variants recovered from supernatants was measured by ELISA with ColE7 as immobilized antigen. All data were normalized to binding activity measured for aglycosylated YebF-Im7 lacking a sequon (wt), such that values greater than 1 (denoted by dashed red line) indicate enhanced binding activity relative to wt Im7. Dashed boxes correspond to two regions (Region 1: residues 23-33; Region 2: residues 58-69) that have many variants with increased activity. Data are average of three biological replicates and error bars represent standard deviation of the mean. (e) DSF analysis of 15 most active YebF-Im7 variants with and without glycosylation. *T*_m_ calculated as midpoint of thermal transition between native and unfolded states. Dashed line indicates *T*_m_ for wt YebF-Im7 (38.6 ± 1.0 °C). Black bars are average of three independent replicates with error bars reported as standard error of the mean.

To exhaustively explore glycosylation sequence space, we constructed all possible individual Im7 sequon variants (80 in total) using the multiplex PCR primer pairs to introduce DQNAT sequons at every position of the protein. A strikingly large number (78 out of 80) of these variants were found to be glycosylated, many with an efficiency that was at or near 100% as estimated from densitometry of the anti-His blot (**Fig. 3c** and **Supplementary Fig. 3**). Because glycosylation by *Cj*PglB can occur both before and after protein folding is completed (**Supplementary Fig. 4**) (32, 33), the secondary and tertiary structure around a glycosylation site is likely to have a direct effect on the extent to which a given site is occupied. Indeed, it has been observed that sequons located in structurally defined regions of folded acceptor proteins are poorly glycosylated and that partial unfolding is required to increase glycosylation efficiency at these sites (33, 34). To determine if the structural context for any of the Im7 sequon variants was a determinant for the timing and efficiency of glycosylation, we performed *in vitro*, cell-free glycosylation reactions in which already folded but yet-to-be glycosylated YebF-Im7 proteins derived from culture supernatants were incubated with purified *Cj*PglB and glycan donor. Remarkably, there was near perfect agreement between the cell-free and cell-based glycosylation results, with nearly all of the purified Im7 variants undergoing highly efficient glycosylation that was at or near 100% with few exceptions (**Supplementary Fig. 3**). The observation that so many Im7 variants were efficiently glycosylated *in vitro* by the *Cj*PglB enzyme (*i.e.*, after folding had been completed) indicates that each sequon was located in either a structurally compliant position (*e.g.*, flexible and surface-exposed loops) within the folded protein or in a region of the protein that became partially unfolded during the cell-free glycosylation reaction. While broad accessibility is certainly plausible given the small size and simple topology of Im7, we cannot rule out the contribution of conformational destabilizing effects caused by substitution of five-residue stretches of native amino acids in the protein. Regardless of the exact reason, these results indicate that Im7 was extremely tolerant to both cell-based and cell-free installation of *N-*glycans over its entire structure.

### Structural and functional consequences of Im7 glycosylation

To exhaustively determine the effect of glycan attachment on neoglycoprotein properties, we first quantified binding activity of all 80 Im7 sequon variants with and without glycosylation by subjecting each to multiwell enzyme-linked immunosorbent assay (ELISA) using purified ColE7 as immobilized antigen. Native Im7 interacts with ColE7, a 60-kDa bacterial toxin that is cytotoxic in the absence of the cognate Im7 inhibitor (35). With an eye towards multiplexibility, we chose to assay YebF-Im7 fusions directly because: (i) it obviated the need for molecular reformatting of the expression constructs; (ii) the fusions could be isolated as relatively pure species from cell-free supernatants, bypassing the need for extensive purification; and (iii) the introduction of the small YebF domain had no measurable effect on ColE7-binding activity (**Supplementary Fig. 5a**). Whereas nearly two thirds of the YebF-Im7 fusions were either unaffected by glycosylation or rendered inactive by introduction of the DQNAT motif alone, particularly in a contiguous stretch between residues 50-57 of Im7, the remaining one third exhibited significantly altered binding activity that was attributable to the presence of the *N-*glycan (**Fig. 3d**). These glycosylation-induced effects were clearly dependent on the precise location of the modification. Indeed, some of the most striking increases in binding activity for glycosylated variants over their aglycosylated counterparts were observed to occur at the transition between different types of secondary structure (*e.g.*, variants Im7^N33^, Im7^N58^ and Im7^N65^). These results were particularly noteworthy in light of the elevated probability of finding naturally occurring sequons in locations where secondary structure changes (31).

Among the Im7 neoglycoproteins whose activity was most significantly affected both positively and negatively by *N-*glycosylation, the majority were located in two distinct regions covering residues 23–33 and 58–69 (**Fig. 3d**). These regions occurred within the two arms of Im7 (one located in α1–loop12–α2 from residue 19 to 39 and the other in loop23–α3–loop34 from residue 46 to 63) that interact extensively with a continuous region in ColE7 in the crystal structure (**Fig. 3c**) (25). The two interfaces are charge-complementary, and charge interactions are largely responsible for the tight and specific binding between the two proteins; hence, it was not surprising that binding activity was sensitive to *N-*glycan attachment in the vicinity of these interfaces. It should be pointed out that the presence of an *N*-glycan in some of these positions was uniquely modulatory, as substitution of DQNAT alone in these same locations generally had little effect on activity, as evidenced by the comparable ColE7 binding measured for aglycosylated Im7 variants versus wt Im7 (**Supplementary Fig. 5b**).

To determine whether any of the glycosylation-induced increases in binding activity were related to stabilization of the native fold, the most active Im7 neoglycoproteins were subjected to differential scanning fluorimetry (DSF) with SYPRO Orange dye in a real-time PCR instrument. Previous studies showed that melting temperature (*T*_m_) values obtained by DSF correlated well with those determined by circular dichroism (CD) thermal denaturation (36). Here too, we observed excellent agreement between these two methods, which both yielded *T*_m_ values for wt Im7 (~39 °C, **Supplementary Fig. 5c and d**) that agreed with a previously reported value (35). Importantly, the presence of the small YebF domain did not significantly alter the *T*_m_ value for Im7 (**Supplementary Fig. 5d**), consistent with its lack of effect on ColE7-binding activity. We also confirmed that DSF results obtained using YebF-Im7 derived directly from cell-free supernatants were indistinguishable from those obtained with more extensively purified YebF-Im7 (**Supplementary Fig. 5d**). Using DSF, the average *T*_m_ values for glycosylated and aglycosylated versions of each Im7 variant were measured, and the change in unfolding temperature, Δ*T*_m_, was calculated such that a positive Δ*T*_m_ signified an increase in structural order and a reduced conformational flexibility due to appending a glycan. Several of the variants exhibited positive Δ*T*_m_ values, with the largest increases corresponding to glycan installation at N33, N59, N60, N65 and N80 (**Fig. 3e**). Conversely, glycans at N10, N58, and N64 caused the largest decreases in *T*_m_, indicative of glycan-induced protein structural changes that destabilized the protein.

### SSGM of an acceptor protein with more complex topology

We next turned our attention to bovine RNase A. Like Im7, RNase A has been intensely studied from a structure–function standpoint and has been pivotal to understanding many aspects of enzymology, biological chemistry, and protein folding and stability. We chose RNase A because (i) it is a relatively small, basic protein, containing 124 residues but with a more complex topology than Im7, with all major types of secondary structure, namely α-helices, β-sheets, and turns, represented; (ii) the natively glycosylated form of RNase A, namely RNase B, contains a single *N*-linked oligosaccharide at N34 and a crystal structure is available (37); (iii) glycosylation at N34 has no apparent effect on the secondary or tertiary structure (37) but does appear to alter the thermal stability (38) although this is controversial (39); and (iv) RNase A modified with an optimal bacterial sequon at the native N34 glycosylation site (RNase A^N34^) can be glycosylated by *Cj*PglB in both cell-based and cell-free reactions (32, 33). For these reasons, RNase A represented an ideal target for SSGM.

Extracellular secretion of glycosylated YebF-RNase A^N34^ was observed in colony blots and immunoblots (**Fig. 2c**), confirming the compatibility of RNase A with glycoSNAP screening. An SSGM library was created by subjecting YebF-RNase A plasmid DNA to the multiplex inverse PCR method, resulting in sequence coverage of 93% in the pre-selected library as determined by next-generation sequencing (**Supplementary Fig. 2**). CLM24 cells carrying plasmids encoding the requisite *C. jejuni* glycosylation machinery were transformed with the SSGM library and subjected to glycoSNAP screening. A total of ~100 glycosylation-positive colonies were randomly selected from two membranes and subjected to sequencing analysis. Of these, only 50 were non-redundant as many of the sequences were isolated multiple times (*e.g.*, seven times each for RNase A^N41^ and RNase A^N122^; **Fig. 4a**). The sequons of these positive hits were uniformly distributed throughout the primary sequence and found in every type of secondary structural element, akin to the results with Im7. Immunoblot analysis confirmed that all selected clones were glycosylated, and the efficiency for most was at or near 100% as estimated by densitometry analysis of the anti-His blots (**Fig. 4b** and **Supplementary Fig. 6a and b**). We also performed theoretical analysis of each of these RNase A glycosite variants in terms of glycosylation probability using NetNGlyc1.0 (http://www.cbs.dtu.dk/services/NetNGlyc/), a web-based tool that predicts *N*-glycosylation sites in human proteins using artificial neural networks that examine the sequence context of N-X-S/T sequons (40). Interestingly, a total of 18 glycosites, which were predominantly clustered in the C-terminal half of the protein, had a glycosylation probability score below 50% (**Supplementary Fig. 6c**) and thus would be predicted to inefficiently glycosylated, if at all. RNase A^N111^ and RNase A^N122^, in particular, both scored below 30% and yet were both very efficiently glycosylated in cells (and *in vitro*, as discussed below).

**Figure 4.**
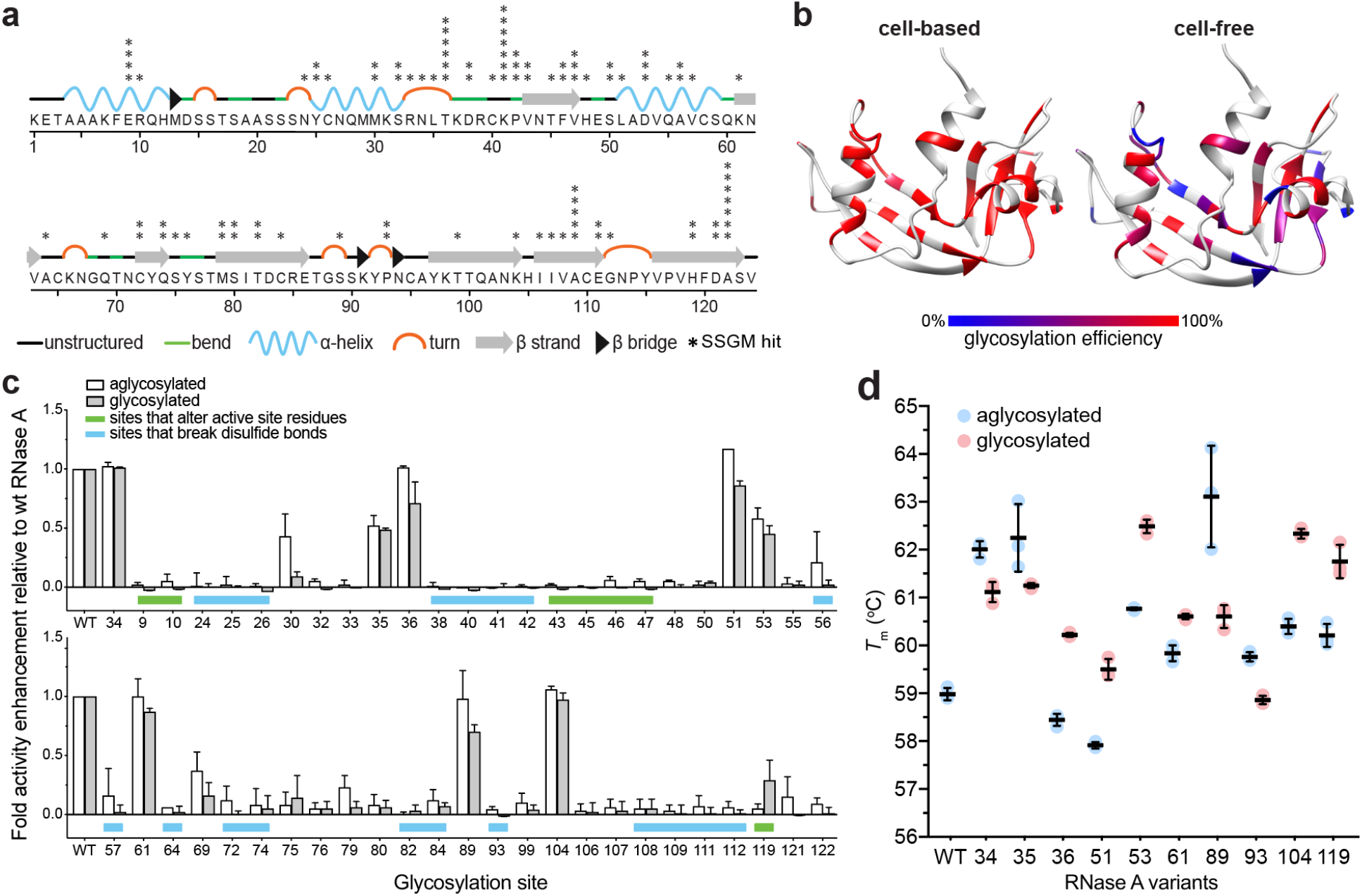
Construction and characterization of RNase A neoglycoprotein libraries. (a) Primary sequence and predicted secondary structure for bovine pancreatic RNase A. Asterisks denote location and frequency of glycosite hits isolated using SSGM. Predicted structures adapted from PDB ID 1RBX. (b) Mapping of cell-based (left) and cell-free (right) glycosylation efficiency onto three-dimensional structure of RNase A. Heatmap analysis of glycosylation efficiency was determined based on densitometric quantification of percent glycosylated (defined as g1/[g0+g1] ratio) for each neoglycoprotein in anti-His immunoblot. (c) Enzymatic activity of glycosylated (gray bars) and aglycosylated (white bars) RNase A variants recovered from culture supernatants. All data were normalized to binding activity measured for aglycosylated YebF-RNase A lacking a sequon (wt). Data are average of three biological replicates and error bars represent standard deviation of the mean. (d) DSF analysis of YebF-RNase A variants with and without glycosylation. *T*_m_ was calculated as midpoint of thermal transition between native and unfolded states. Dashed line indicates *T*_m_ for wt YebF-RNase A (59.0 ± 0.1 °C). Black bars are average of three independent replicates with error bars reported as standard error of the mean.

To investigate whether the structural context of the sequon impacted the possible timing of PglB-mediated glycan installation, we performed *in vitro*, cell-free glycosylation of folded RNase A variants. While some variants were glycosylated equally well in cell-based and cell-free reactions (*e.g.*, RNase A^N46^ and RNase A^N64^), an unexpectedly large number showed significantly lower levels of glycosylation under cell-free conditions (**Fig. 4b** and **Supplementary Fig. 6a and b**). Most notably among these were variants N34, N35, N36, N43, N51, N61, N69, N72, N80, N89, and N104, which were all efficiently glycosylated in cells but underwent little or no detectable glycosylation *in vitro*. These sequons occur at locations that were likely to be accessible to the OST during translation/translocation when the proteins are unfolded but became inaccessible after the protein completed folding. Indeed, the native *N*-glycosylation site at N34 is located in a structured domain, suggesting that the poor cell-free glycosylation at this specific location (and perhaps also at the nearby N36 and N43 sites) was due to sequon inaccessibility in the folded state. Such folding-dependent recognition of this site has been observed previously (32, 33) and, together with the results presented here, supports a model whereby cell-based glycosylation of these particular sequons involves glycan installation prior to folding, either co- or post-translocationally (**Supplementary Fig. 4**).

To determine the consequences of glycosylation at the 50 unique sites, the ability of glycosylated and aglycosylated versions of each sequon variant to catalyze the hydrolysis of the phosphodiester bonds in RNA was evaluated. While the addition of YebF had little to no effect on RNase A activity (**Supplementary Fig. 7a**), more than half of the RNase A variants were inactivated by substitution of the DQNAT sequon (**Fig. 4c**). To determine if this might be due to the substitution of five residues in the target protein, a requirement for optimal recognition by *Cj*PglB (41), we mutated RNase A more conservatively at a select number of sites. Specifically, we generated minimal sequons (D-X-N-X-T/S or X-X-N-X-T/S, where X represents the native amino acid), which in most cases required only 1 or 2 amino acid changes. Each of these mutants was completely inactive except for RNase A^N55^ with a DVNAT sequon, which retained some activity but was still significantly less active than the wt enzyme (**Supplementary Fig. 7b**). Hence, even relatively minor sequence perturbations at these positions, in addition to the less subtle substitution with DQNAT, were all capable of inactivating RNase A. More careful inspection revealed that the majority of variants with little to no activity corresponded to the substitution of sequons in locations that would be predicted to disrupt catalytically important residues or disulfide bonds (**Fig. 4c** and **Supplementary Results**).

Among the RNase A neoglycoproteins that retained function, only eight (sequons at N34, N35, N36, N51, N53, N61, N89, and N104) showed activity that was on par (>50%) with wt RNase A but none were more active than their aglycosylated counterpart (**Fig. 4c**). In the case of RNase A^N119^, introduction of the DQNAT sequence completely abrogated catalytic activity, consistent with previous findings that the relative activity of an H119N mutant was reduced to less than 1% of wt RNase A, with *k*_cat_/*K*_M_ values reduced by 100- to 1000-fold depending on the substrate used (42). Despite the importance of this residue for catalysis, glycosylation at this position partially restored enzymatic activity, indicating an *N*-glycan-dependent gain-of-function.

To determine whether glycosylation impacted stability, we again used DSF to analyze the most active RNase A neoglycoproteins along with RNase A^N93^, which was randomly chosen as a representative inactive variant. The measured *T*_m_ values for wt YebF-RNase A and its unfused counterpart were both ~59 °C (**Supplementary Fig. 7c**), in close agreement with previous findings (39), while the *T*_m_ values for all the YebF-RNase A variants spanned a range from 58–63 °C (**Fig. 4d**). Most exhibited positive Δ*T*_m_ values compared to their aglycosylated counterpart, including the RNase A^N119^ variant, suggesting that the restoration of activity caused by glycan attachment at N119 also served to stabilize the protein. In contrast, RNase A^N89^ and RNase A^N93^ exhibited large negative Δ*T*_m_ values that coincided with slightly weakened activity due to glycan attachment in the case of N89 and complete inactivation in the case of N93.

### Investigation of IgG variable domain glycosylation using SSGM

We next investigated antibody variable domain glycosylation, a phenomenon that is observed for ~15% of serum IgGs and contributes to diversification of the B-cell antibody repertoire (6). Although glycan installation within the variable domains of Fab arms has been long known, the rules governing the selection of *N*-glycosylation sites in Fab domains that emerge during somatic hypermutation and the functional consequences of the attached glycans remain poorly understood. To systematically investigate this phenomenon using SSGM, the two variable domains, V_H_ and V_L_, from the human anti-HER2 monoclonal antibody were joined by a flexible linker to form scFv-HER2 that was subsequently modified at its N-terminus with YebF and at its C-terminus with a DQNAT motif. Extracellular secretion of glycosylated YebF-scFv-HER2^DQNAT^ was observed in colony blots and immunoblots (**Fig. 5a**), confirming the compatibility of scFv-HER2 with glycoSNAP screening. Because variable domain glycosylation is subject to selection mechanisms that depend on the nature of the antigen (6), we modified the SSGM strategy to enable dual screening of glycosylation and antigen-binding activity by labeling colonies with SBA lectin and the extracellular domain (residues 1-652) of human HER2 (HER2-ED), which was avidly bound by scFv-HER2^DQNAT^ fused to YebF (**Supplementary Fig. 8a**). In this way, two-color screening could be used to identify colonies that were positive both for glycosylation and for antigen binding, as demonstrated with the YebF-scFv-HER2^DQNAT^ construct (**Fig. 5a**). Next, we constructed and screened an SSGM library, after which two-color glycoSNAP screening was performed with CLM24 cells carrying plasmids encoding the library and the *C. jejuni* glycosylation machinery. A total of ~60 dual-positive hits were isolated from membranes, of which 21 were determined to be non-redundant (*e.g.*, N58 in V_L_ and N42 in V_H_ were each isolated 12 times) (**Fig. 5b**) and subsequently confirmed for extent of glycosylation by immunoblot and densitometry analysis (**Supplementary Fig. 8b and c**). The sequons of these hits were sparsely distributed throughout the primary sequence, with a large proportion clustering just after the second and third complementarity-determining regions (CDRs) of the V_L_ domain and also in the flexible linker, indicating a clear selection bias for specific sites that tolerated glycosylation without interfering with binding function. Interestingly, a few of the identified sequons occurred in CDR2 of the V_L_ domain and CDR1 and CDR2 of the V_H_ domain, consistent with naturally occurring IgG repertoires in which *N*-glycosites are found preferentially in the CDRs (6).

**Figure 5.**
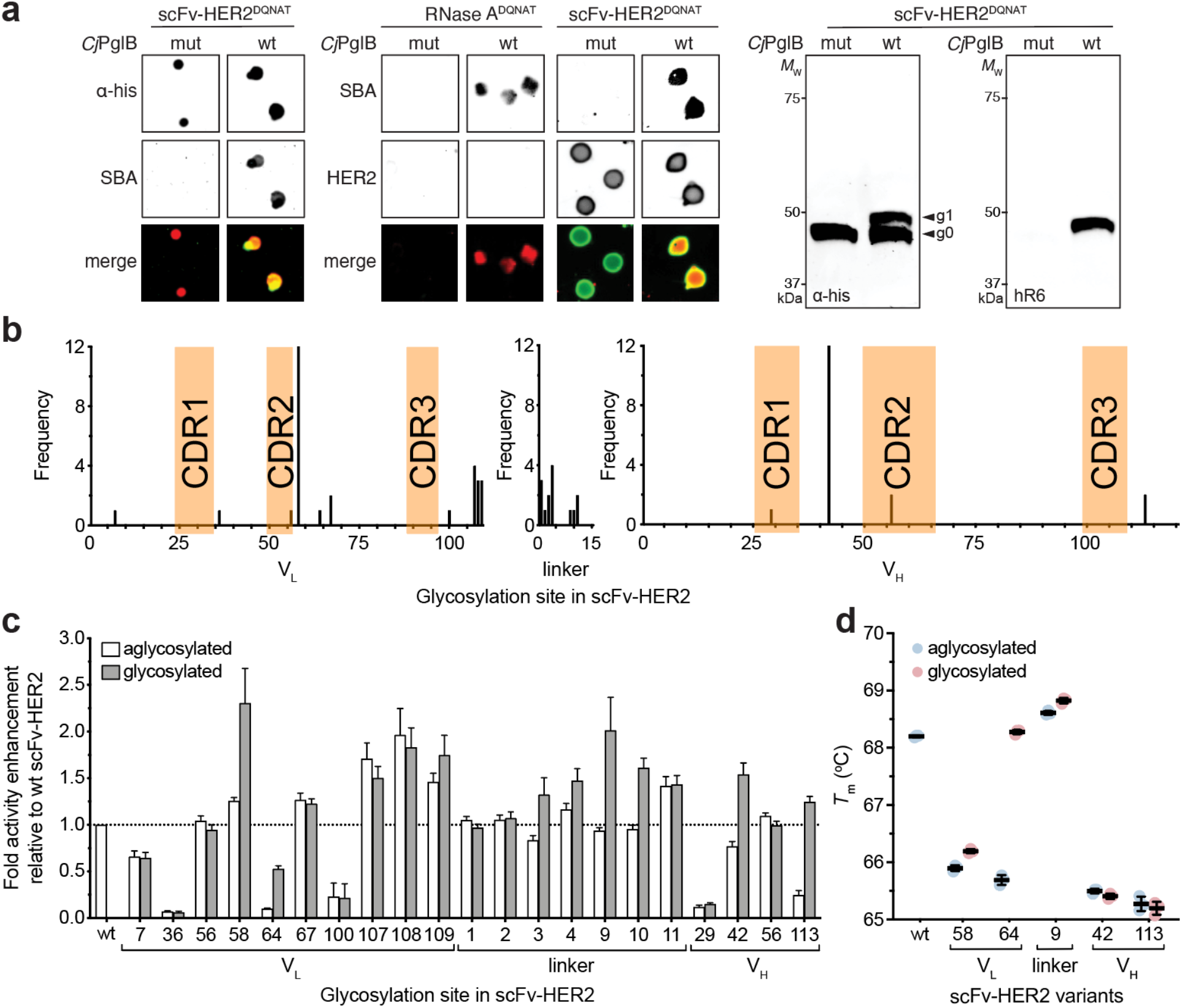
Construction and characterization of scFv-HER2 neoglycoprotein libraries. (a) Immunoblot analysis of acceptor proteins in colony secretions (left and middle) and periplasmic fractions (right) derived from *E. coli* CLM24 carrying plasmids encoding scFv-HER2^DQNAT^ and requisite *N*-glycosylation machinery with either wild-type *Cj*PglB (wt) or an inactive mutant (mut). Blots were probed with anti-polyhistidine antibody (α-His) to detect acceptor protein, SBA or hR6 serum to detect the glycan, and HER2-ED to detect antibody binding. Bottom color panels depict overlay of α-His and SBA blots or SBA and HER2 blots (merge). Arrows denote aglycosylated (g0) and singly glycosylated (g1) forms of scFv-HER2^DQNAT^. Molecular weight (*M*_W_) markers are indicated at left. Results are representative of at least three biological replicates. (b) Frequency and position of *N*-glycosylation sites in scFv-HER2^DQNAT^ glycovariants isolated using SSGM. (c) Binding activity of glycosylated (gray bars) and aglycosylated (white bars) scFv-HER2^DQNAT^ variants as measured by ELISA with HER2-ED as immobilized antigen. All data were normalized to binding activity measured for aglycosylated scFv-HER2 lacking a sequon (wt), such that values greater than 1 (denoted by dashed line) indicate enhanced binding activity relative to wt scFv-HER2. Data are average of three biological replicates and error bars represent standard deviation of the mean. (d) DSF analysis of YebF-scFv-HER2 variants with and without glycosylation. *T*_m_ was calculated as midpoint of thermal transition between native and unfolded states. Dashed line indicates *T*_m_ for wt YebF-scFv-HER2 (68.2 ± 0.1 ^°^C). Black bars are average of three independent replicates with error bars reported as standard error of the mean.

In terms of function, all 21 scFv-HER2 hits exhibited HER2-ED binding activity above background (**Fig. 5c**), which was expected given that the screening process was adapted to include antigen binding. Importantly, nine of these neoglycoproteins (N58, N64, and N109 in V_L_; N3, N4, N9, N10 in linker; N42 and N113 in V_H_) exhibited increased binding compared to their aglycosylated counterpart, and most of these were also more active than the parental scFv-HER2. For the five clones exhibiting the greatest increase in activity due to glycosylation, we measured *T*_m_ values and found that in general glycan attachment did not affect stability (**Fig. 5d**). However, the one exception was N64 V_L_, which experienced a 2.6 °C increase in *T*_m_ due to the addition of the *N-*glycan. Overall, these results are in agreement with several previous studies showing that variable region glycans contribute to antibody binding characteristics and stability in a manner that depends on the precise location of the glycan (6, 43) and suggest that glycosylation in this region may be a useful strategy for fine-tuning the performance of IgG antibodies and their engineered derivatives.

### Computational analysis of neoglycoproteins

To test whether protein-structure analyses could explain the observed effects of sequon substitution and glycosylation, we modeled the sequon-substituted variants, with and without glycosylation, and calculated simple geometric measures (secondary structure, burial, distance to the binding site, and surface area) as well as Rosetta energy estimates (stability and interface score) for each. Unfortunately, none of these factors were found to correlate significantly with the activity or stability of the Im7 or RNase A neoglycoproteins (**Supplementary Figs. 9-13**; and **Supplementary Results**). It should be noted that these metrics may be less useful for RNase A because the activities are primarily explained by the disruption of the active site and the disulfide bonds, which are not captured in these metrics. We also generated ensembles of glycan conformations for several Im7 neoglycoproteins, including several with increased (Im7^N30^, Im7^N49^, and Im7^N58^) and one with decreased (Im7^N31^) activity due to glycosylation, in the context of binding to E7. These ensembles revealed that: (i) the glycan and the bound protein often interact to change the binding activity positively or negatively; and (ii) enhanced binding appears to be mediated by multiple low-energy glycan conformations making favorable interactions with E7 (**Supplementary Fig. 14**; and **Supplementary Results**).

Next, we compared the experimental binding activity for scFv-HER2 with multiple geometric and Rosetta metrics. Unlike Im7 or RNase A, scFv-HER2 activity correlated with many of our metrics. First, sequon burial reduces the binding affinity of scFv-HER2 for its antigen both in the glycosylated (*R*^2^ = 0.43) and aglycosylated (*R*^2^ = 0.21) states (**Fig. 6a** and **Supplementary Fig. 9b**, respectively). Similarly, the closer the sequon was to the paratope, the greater the likelihood of reduced activity for the glycosylated (*R*^2^ = 0.23) and aglycosylated (*R*^2^ = 0.20) variants (**Fig. 6b** and **Supplementary Fig. 9c**, respectively). The buried surface area also correlated with the activity of the glycosylated variant (*R*^2^ = 0.19, **Supplementary Fig. 10e**). The strongest predictors, however, were the Rosetta scores. For the glycosylated state, the activity correlated with both the total Rosetta score (*R*^2^ = 0.49, **Fig. 6c**) and the interface score (*R*^2^ = 0.63, **Supplementary Fig. 10g**). The aglycosylated antibody–antigen complex total score correlated with experimental binding activity (*R*^2^ = 0.49, **Supplementary Fig. 9f**). These Rosetta scores were primarily driven by the van der Waals complementarity and to a lesser extent electrostatics (**Supplementary Figs. 11** and **12**).

**Figure 6:**
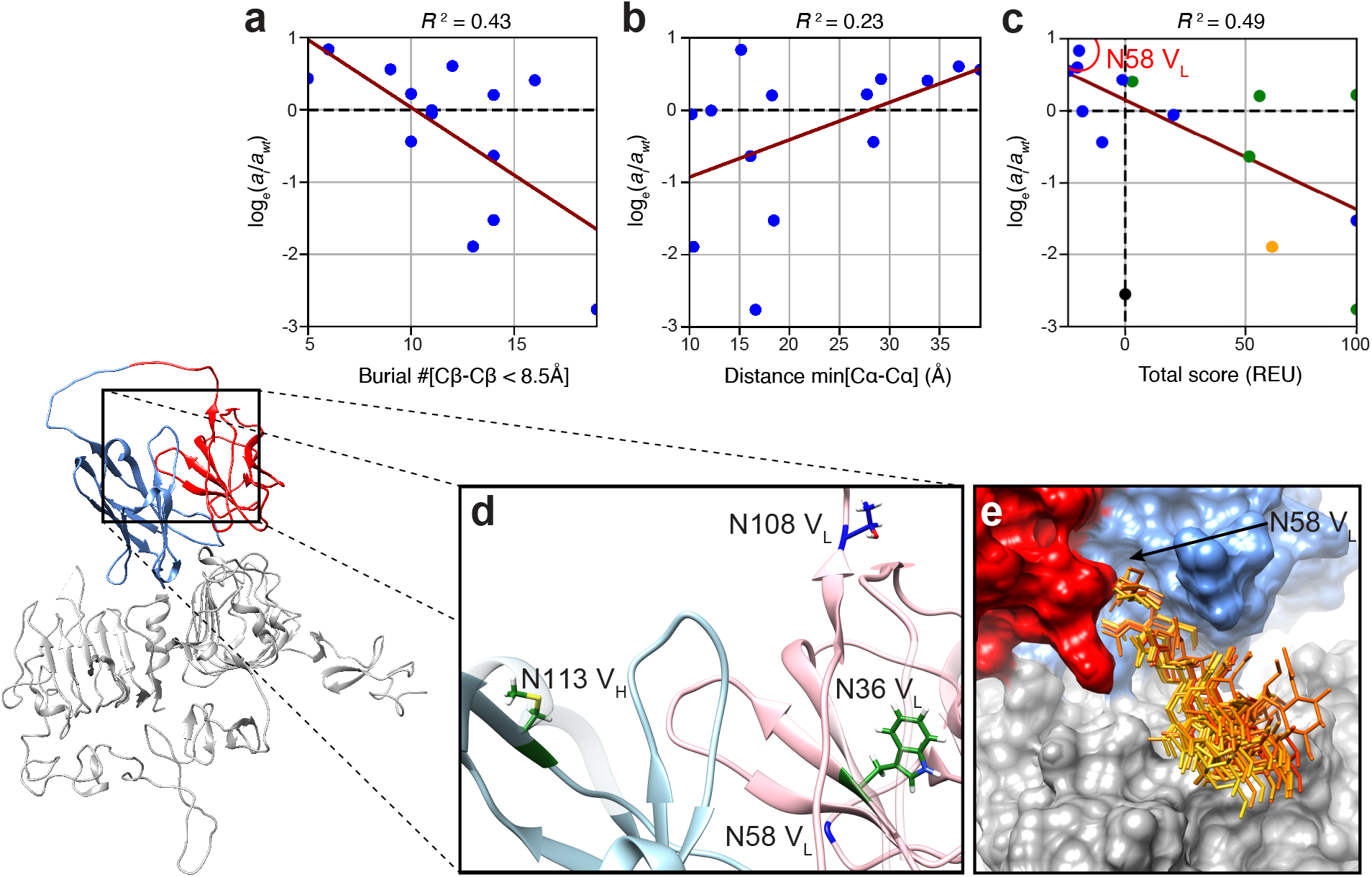
Computational analysis of scFv-HER2 neoglycovariants. The structure of scFv-HER2 V_L_ (red) and V_H_ (blue) domains in complex with HER2 protein (gray) is shown at bottom left. Regression analyses of log activity ratio (glycosylated / wild-type) versus (a) burial of sequon substitution site, (b) distance of closest HER2 residue from the sequon substitution site, and (c) total Rosetta score. In all three panels, the dark red lines are the respective regression lines. Colors of dots in (c) show the respective secondary structure of the sequon substitution site. Orange, green, and blue correspond to α-helix, β-strand, and loop regions, respectively. N58 V_L_ (red circle) has the highest glycosylated binding activity increase and is discussed in the text. (d) Wild-type representation of sites used for analysis of sequon substitution (36 V_L_, 108 V_L_, and 113 V_H_) and glycosylation (58 V_L_). Side-chain colors reflect their respective secondary structures. (e) Glycan arrangement (orange sticks) from eight low energy conformations of glycosylated N58 V_L_ variant of scFv-HER2, revealing possible glycan-HER2 interaction responsible for binding activity improvement.

For the aglycosylated activities, we selected three variants for deeper analysis: two variants that had low binding activity and a poor Rosetta score (N36 V_L_, N113 V_H_; black circles in **Supplementary Fig. 11a**) and one variant with high activity and a favorable Rosetta score (N108 V_L_; red circle in **Supplementary Fig. 11a**). Both N36 V_L_ and N113 V_H_ sites are situated on β-strands in compact regions of the anti-HER2 antibody on the side opposite the antigen-binding site (**Fig. 6d**, green sticks). The reduced stability arises from the steric clash of substituting a sequon inside (or near) a close-packed region of the protein (Rosetta terms for steric clashes (vdW_rep) of 90.2 and 79.8 Rosetta energy units (REU) for the N36 V_L_ and N113 V_H_, respectively). When glycosylated, the clashes worsen in the Rosetta models, corresponding to low activity (black circles in **Supplementary Fig. 11a**). On the other hand, site N108 V_L_ is located at the C-terminal end of V_H_ (**Fig. 6d**, blue sticks). Sequon substitution had a relatively small effect on the electrostatic interactions (−6.2 REU) and a greater effect on the repulsive van der Waals terms (−28.0 REU), indicating that new side chains are acceptable in less compact regions. A similar outcome was reported following substitution mutation of a human monoclonal antibody (44).

To understand how *N-*glycosylation was able to improve binding activity, we selected mutant N58 V_L_ because the aglycosylated variant was 26% more active than the wt scFv-HER2 and glycan addition improves the binding an additional 1.8-fold. Residue N58 V_L_ resides in the turn between strands 1 and 2 (**Fig. 6d**, blue backbone). From Rosetta-generated glycosylated structures, the low-energy states showed interfacial contacts between the glycan and the surface residues of HER2 (**Fig. 6e**), improving both the total Rosetta score and the interface score (red circle in **Fig. 6c** and **Supplementary Fig. 10g**) and explaining the binding activity improvement as resulting from favorable glycan-antigen contacts.

## Discussion

In this study, we developed a new protein engineering workflow called SSGM for constructing large neoglycoprotein libraries of virtually any POI and characterizing the consequences of glycan installation. The utility and flexibility of this technique was demonstrated using three structurally and functionally diverse acceptor proteins: bacterial Im7, bovine RNase A, and human scFv-HER2. Specifically, each of these proteins was subjected to a systematic “sequon walking” procedure that enabled creation of synthetic gene libraries in which *N-*glycosylation sites (the majority of which were naïve) were introduced at every possible position of the POI. Upon screening these libraries using glycoSNAP (21), numerous positions in each protein were found to be efficiently *N-* glycosylated. While extended regions and loops tended to be more receptive to glycosylation, all types of secondary structure were found to be glycosylated, consistent with the observation that naturally occurring *N-*glycans also exist on all forms of secondary structure (31). For RNase A, in particular, a significant number of the efficiently glycosylated sites (18/50) were predicted to have very low glycosylation potential, highlighting the need for large-scale experimental studies of glycosylation, such as described here, that can be used to help refine predictive tools. To this end, higher throughput techniques that leverage mass spectrometry for quantitatively resolving glycosylation efficiency (45, 46) could enable further refinement of the method in the future.

The studies performed here also provided insight on the possible timing and impact of glycosylation with respect to the folding process. For instance, Im7 tolerated a glycan at almost every position, even when the target asparagine side chain pointed inward and was considered buried (*e.g.*, positions N7, N68, and N76). Because these buried positions physically cannot be glycosylated by PglB when the target protein is in the folded state, they must either be glycosylated co-translationally or during a fluctuation to a partially unfolded state that provides access to that site. Then, after glycosylation, because Im7 presumably cannot fold back into the native structure, it must adopt a different conformation to accommodate the newly added glycan, which would be feasible in light of the fact that Im7 is very flexible (47). In the case of RNase A, several sites were identified (*e.g.*, N34, N36) that could be efficiently glycosylated in cells but underwent little to no glycosylation *in vitro* (in the already folded state), providing clear evidence for glycan installation prior to folding and in a manner that may resemble the co-translocational process in mammalian cells (48). The overall less efficient glycosylation seen for many RNase A variants was also consistent with the protein adopting a more stable folded structure compared to Im7 and providing less accessibility to buried sites.

In addition to uncovering glycosylatable sites, the SSGM workflow also allowed the effects of these site-directed glycan “mutations” to be probed for their contribution to the biological and biophysical properties of each POI. In this way, SSGM is conceptually analogous to combinatorial alanine-scanning mutagenesis, which allows systematic determination of the importance of individual amino acids to protein structure and function (49–51). Consistent with the known modulatory effects of *N-*glycans (4, 5), many of the neoglycoprotein derivatives of Im7, RNase A and scFv-HER2 exhibited detectably altered stability and activity that resulted from covalent attachment of *N-*glycans at precise locations in the protein backbone. For example, installing *N-*glycans in the center of α-helices negatively affected activity (*e.g.*, positions 19, 42, 72 in Im7) whereas those installed at the transition between different types of secondary structure and at turns between motifs promoted enhanced activity and, in some cases, stability (*e.g.*, positions 33, 49, 58, 59, 60, 61, 65, 67, 68, 69, 78, 80 in Im7). These findings generally agreed with the folding and stability effects contributed by attachment of a GlcNAc_2_ disaccharide to discrete locations in Im7 (20) and also provide clues for why natural *N*-glycosylation sites occur with elevated frequency in turns and bends and especially at points of change in secondary structure and with low frequency within ordered helices (31). Despite the overall agreement with previous studies, a few notable differences emerged. For example, in our hands, Im7 glycosylated at position 27 with the GalNAc_5_(Glc)GlcNAc heptasaccharide was more active but equally stable as its aglycosylated counterpart, whereas an EPL-derived Im7 modified with chitobiose at residue 27 was significantly more stable than unmodified Im7 (note that activity data was not reported) (20). Likewise, RNase A^N34^ glycosylated with GalNAc_5_(Glc)GlcNAc exhibited activity that was nearly identical to that of aglycosylated RNase A^N34^ (and wt RNase A), whereas the attachment of oligomannose glycans at N34 was previously observed to reduce activity by more than threefold (52). The notion that discrete glycan structures attached to the same site in a protein can have disparate effects is not unprecedented, having been documented for other glycoproteins (53) (54). Thus, in the future, it will of interest to extend SSGM for use with alternative glycan structures, including for example Man_3_GlcNAc_2_ or other human *N-* and *O-*linked glycans that have been engineered in *E. coli* (32, 55, 56), so that the consequences of varying glycan structures at discrete locations can be systematically investigated.

The fact that *N-*glycan attachment significantly increased the binding activity of several glycosite variants of Im7 and scFv-HER2 suggests that SSGM may become a useful tool for adding *N-*glycans to naïve sites in proteins for tuning their biological and biophysical properties. The discovery of such sites was accelerated by the ability of SSGM to furnish an unprecedentedly large number of intact neoglycoproteins (a total of 151 in this study alone), for which the effects of *N*-glycan installation can be readily catalogued using multiplexable assays for protein structure and activity as we showed here. While no definitive rules regarding the effects of glycosylation were revealed here, we anticipate that sequon walking on a larger, even proteome-wide, scale could provide access to datasets that might allow the effects of glycosylation to be more widely generalized and perhaps even predicted. Nonetheless, computational analysis indicated that interactions between the glycan and the bound protein can alter binding activity (positively or negatively) and that enhanced binding likely arises from low-energy glycan conformations making favorable interactions with the binding partner. For example, the Im7^N58^ variant that underwent the largest increase in binding activity upon glycosylation also acquired new contacts with its binding partner, E7, through the glycan, which strengthened binding activity 3.5-fold. Likewise, for the scFv-HER2 mutant N58 V_L_, which exhibited measurably higher antigen-binding activity compared to parental scFv-HER2, the heptameric glycan created new contacts between scFv-HER2 and HER2-ED and buried more surface area upon binding. Thus, even though part of the enhanced binding was from the sequon substitution alone, perhaps from the additional contacts of the long Q57 side chain or from a stabilizing effect of the sequon on the CDR L2 loop (residues 51-57 in V_L_), most of the effect was from the *N-*glycan itself. Importantly, this observation was in line with previous findings that glycans attached near (but not within) the antigen-binding site can increase affinity (57). Taken together, our findings suggest that SSGM could be used to rapidly identify naïve sites along a protein backbone for strategic placement of *N-*glycans that substantially enhance the biological and/or biophysical properties of the resulting neoglycoprotein.

## Materials and Methods

### Strains and culture conditions

*E. coli* strain DH5α was used for all molecular biology, including plasmid construction, site-directed mutagenesis, and SSGM library construction. BL21(DE3) was used to purify ColE7 that was used to measure Im7 binding activity in ELISA format. All glycosylation studies were performed using *E. coli* strain CLM24 (58), which was initially grown at 37 °C in Luria–Bertani (LB) medium containing appropriate antibiotics at the following concentrations: 20 μg/mL chloramphenicol (Cm), 100 μg/mL trimethoprim (Tmp), and 50 μg/mL spectinomycin (Spec). When cells reached mid-log phase, protein expression was induced by adding 0.1 mM isopropyl-β-D-thiogalactoside (IPTG) and 0.2% (v/v) L-arabinose, after which cells were grown at 30 °C for 16–20 h.

### Plasmid construction

For expression of the glycosylation machinery, plasmid pMAF10 encoding *Cj*PglB (58) along with either plasmid pMW07-pglΔB (21) or pMW07-pglΔBCDEF were used. The latter plasmid was constructed by deleting the *pglCDEF* genes from plasmid pMW07-pglΔB (21), resulting in a modified *C. jejuni* glycan biosynthesis pathway that excluded the *pglB* gene encoding the OST and also the *pglCDEF* genes encoding enzymes for synthesis and transfer of bacillosamine as described previously (27). This deletion can be complemented by *E. coli* WecA, a sugar-phosphate transferase that transfers GlcNAc phosphate to undecaprenol phosphate, therefore initiating LLO biosynthesis. It should be noted that while pMW07-pglΔB encodes the bacillosamine-related *pglCDEF* genes, we did not detect the presence of bacillosamine in any of the glycoforms produced by cells carrying this plasmid. A derivative of pMAF10 that encoded a catalytically inactive version of *Cj*PglB carrying two active-site mutations (D54N and E316Q) (21) was used as a negative control. The plasmids pTrc99S-YebF-Im7 and pTrc99S-YebF-Im7^DQNAT^ were constructed by inserting cloning cassettes YebF^N24L^-XbaI-Im7-SalI-FLAG-6xHis and YebF^N24L^-XbaI-Im7-BamHI-DQNAT-SalI-FLAG-6xHis, respectively, into the SacI and HindIII sites of pTrc99S (59). The genes encoding RNase A and scFv-HER2 were PCR amplified from plasmids pTrc-ssDsbA-RNaseA (21) and pMAZ360–cIgG–Herceptin (60), respectively, and cloned into the cassette between XbaI and SalI sites in pTrc99S-YebF-Im7 or XbaI and BamHI sites in pTrc99S-YebF-Im7^DQNAT^, replacing Im7 and Im7^DQNAT^, respectively. The pTrc-spDsbA-POI plasmids (where POI corresponds to each of the proteins of interest, namely Im7, RNase A, and scFv-HER2) were cloned by one-step PCR integration of primers encoding the *E. coli* DsbA signal peptide (spDsbA) into each pTrc99S-YebF-POI plasmid as templates followed by Gibson assembly. PCR products were subjected to DpnI digestion to remove parental plasmid. The resulting PCR products were assembled by Gibson assembly and used to transform *E. coli* cells to obtain the desired plasmids. Plasmid pET28-ColE7 (H569A) was constructed by inserting DNA encoding the ColE7 H569A variant (61) bearing a C-terminal 6×His tag (Integrated DNA Technologies) into the NcoI and SalI sites of pET28a. All plasmids were confirmed by DNA sequencing at the Biotechnology Resource Center of the Cornell Institute of Biotechnology.

### SSGM library construction

SSGM mutagenesis libraries were constructed by multiplex inverse PCR (30) followed by T4 ligation. Each of the pTrc99S-YebF-POI plasmids was used as template for PCR amplification using primer sets specifically designed such that the DNA sequence 5′-GAT CAG AAT GCG ACC-3′ was included in the 5′ end of every forward primer to enable substitution of the adjacent five amino acids with DQNAT. Prior to PCR, the forward primers were phosphorylated using T4 polynucleotide kinase (New England Biolabs) to facilitate T4 ligation later. PCR reactions were performed using Phusion polymerase (New England Biolabs), and the PCR products were gel-purified from the product mixtures to eliminate non-specific PCR products. The resulting PCR products were self-assembled using T4 ligase (New England Biolabs) to obtain the desired SSGM plasmid libraries, which were subsequently used to transform highly competent DH5α cells and then isolated using a QIAprep Spin Miniprep Kit (Qiagen) according to manufacturer’s instructions. For next-generation sequencing, see **Supplementary Methods**.

### GlycoSNAP assay

Screening of SSGM libraries was performed using glycoSNAP as described previously (21). Briefly, *E. coli* strain CLM24 carrying pMW07-pglΔB and pMAF10 was transformed with corresponding SSGM library plasmids, and the resulting transformants were grown on 150-mm LB-agar plates containing 20 μg/mL Cm, 100 μg/mL Tmp, and 50 μg/mL Spec overnight at 37 °C. The second day, nitrocellulose transfer membranes were cut to fit 150-mm plates and pre-wet with sterile phosphate-buffered saline (PBS) before placement onto LB-agar plates containing 20 μg/mL Cm, 100 μg/mL Tmp, 50 μg/mL Spec, 0.1 mM IPTG, and 0.2% (w/v) L-arabinose. Library transformants were replicated onto 142-mm nitrocellulose membrane filters (Whatman, 0.45 μm), which were then placed colony-side-up on transfer membranes and incubated at 30 °C for 16 h. The nitrocellulose transfer membranes were washed in Tris-buffered saline (TBS) for 10 min, blocked in 5% bovine serum albumin for 30 min and probed for 1 h with fluorescein-labeled SBA (Vector Laboratories, catalog # FL-1011) and Alexa Fluor 647^®^ (AF647)-conjugated anti-His antibody (R&D Systems, catalog # IC0501R) or HER2-ED (R&D Systems, catalog # 10126-ER) that was conjugated with Alexa Fluor 647^™^ (AF647) (Thermo Fisher Scientific, catalog # A37573) following the manufacturer’s instructions. All positive hits were re-streaked onto fresh LB-agar plates containing 20 μg/mL Cm, 100 μg/mL Tmp, 50 μg/mL Spec, and grown overnight at 37 °C. Individual colonies were grown in liquid culture and subjected to DNA sequencing to confirm the location of glycosites and to protein glycosylation analysis as described below.

### Protein isolation

For Western blot analysis and protein activity assays, cell-free culture supernatants were generated by subjecting 1.5 mL of cells that had been induced for 16 h to centrifugation at 13,4000 × g at 4 °C for 2 min. Periplasmic fractions were generated by subjecting 3 mL of 16-h-induced cultures to centrifugation at 13,400 × g for 2 min. The resulting pellets were resuspended in 300 μL of 0.4 M arginine and incubated at 4 °C for 1 h with gentle shaking. After centrifugation at 13,400 × g for 2 min, the supernatant containing periplasmic extracts was collected. For stability assays, YebF-Im7, YebF-RNase A, and YebF-scFv-HER2 variants were purified from supernatant fractions and soluble lysate fractions. To prepare the latter, cells expressing YebF-RNase A variants were harvested by centrifugation at 6000 × g at 4 °C for 20 min and the pellets were resuspended in PBS buffer supplemented with 10-mM imidazole followed by cell lysis using a Emulsiflex-C5 Homogenizer (Avestin) at 16,000–18,000 psi. The resulting lysate was clarified by centrifugation at 15,000 × g for 30 min at 4 °C to collect the soluble fraction. All soluble fractions, or supernatant fractions supplemented with 10-mM imidazole, were then applied twice to a gravity flow column loaded with Ni-NTA resin at room temperature and washed with PBS containing 20-mM imidazole until the concentration was lower than 0.1 mg/mL. Proteins were eluted in 2.5 mL of PBS with 250 mM imidazole. The eluted proteins were desalted using PD10 Desalting Columns (GE Healthcare) and stored at 4 °C.

To produce ColE7 for ELISA experiments, an overnight culture BL21(DE3) cells carrying plasmid pET28a-ColE7 (H569A) was used to inoculate 1 L of LB supplemented with 50 μg/mL kanamycin. Cells were grown at 37 °C until mid-log phase and then were induced with 0.1 mM IPTG for 16 h at 16 °C before being harvested. Following centrifugation at 10,000× g, pellets were resuspended in PBS buffer supplemented with 10-mM imidazole and lysed at 16,000–18,000 psi using an Emulsiflex-C5 homogenizer (Avestin). The lysate was clarified by centrifugation at 15,000 × g for 30 min at 4 °C and the collected soluble fraction was mixed with Ni-NTA resin for 2 h at 4 °C. The mixture was then applied to a gravity flow column and washed with 5 column volumes of PBS containing 20 mM imidazole. Proteins were eluted in 4 column volumes of PBS with 250-mM imidazole. The eluted protein was desalted and concentrated to 5 mg/mL in PBS buffer using Ultra Centrifugal Filters with 10-kDa molecular weight cut-off (Amicon^®^) and stored at 4 °C.

### Western blotting analysis

Supernatant or periplasmic fractions were diluted 3:1 in 4× Laemmli sample buffer (Bio-Rad) and were boiled at 100 °C for 10 min. The treated samples were subjected to SDS-polyacrylamide gel electrophoresis on 10% Mini-PROTEAN^®^ TGX™ Precast Protein Gels (Bio-Rad). The separated protein samples were then transferred to nitrocellulose membranes. Following transfer, the membranes were blocked with 5% milk (w/v) in TBST (TBS, 0.1% Tween 20) and were probed with horseradish peroxidase (HRP) conjugated anti-His antibody (Abcam, catalog # ab1187) or the *C. jejuni* heptasaccharide glycan-specific antiserum hR6 for 1 h. For the latter, goat anti-rabbit IgG (HRP) (Abcam, catalog # ab205718) was used as the secondary antibody to detect hR6 antiserum. After washing three times with TBST for 10 min, the membranes were visualized using a ChemiDoc^™^ MP Imaging System (Bio-Rad). For all methods related to MS analysis of proteins, see the **Supplementary Methods**.

### Cell-free glycosylation

Methods for purification of *C. jejuni* PglB and isolation of LLOs from glycoengineered *E. coli* were described previously (62). *In vitro*, cell-free glycosylation was carried out in 30-μL reactions containing either 20 μL of supernatant fraction containing aglycosylated YebF-Im7 or 20 μL of periplasmic fraction containing YebF-RNase A, 2 μg of purified *Cj*PglB, and 5 μg extracted LLOs in cell-free glycosylation buffer (10-mM HEPES, pH 7.5, 10-mM MnCl2, and 0.1% (w/v) *n*-dodecyl-β-D-maltoside (DDM)). Reaction mixtures were incubated at 30 °C for 16 h and stopped by adding 10 μL of 4× Laemmli sample buffer containing 5% β-mercaptoethanol followed by boiling at 100 °C for 15 min, after which they were subjected to Western blot analysis.

### ELISA

Binding activity for Im7 and scFv-HER2 was determined by standard ELISA. Briefly, Costar 96-well ELISA plates (Corning) were coated overnight at 4 °C with 50 μL of 5 μg/mL purified ColE7 in 0.05-M sodium carbonate buffer (pH 9.6) for Im7 variants and 50 μL of 0.2 μg/mL HER2-ED (Sino Biological, catalog # 10004-HCCH) in PBS buffer for scFv-HER2 variants. After blocking with 5% (w/v) non-fat milk in PBS for 1 h at room temperature, the plates were washed three times with PBST (PBS, 0.05% (v/v) Tween-20) and incubated with serially diluted aglycosylated and glycosylated YebF-Im7 and YebF-scFv-HER2 glycovariants for 1 h at room temperature. After washing three times with PBST, 50 μL of 1:2,500-diluted HRP-conjugated anti-DDDK tag antibody (Abcam, catalog # ab49763) for Im7 variants or 50 μL of 1:5,000-diluted HRP-conjugated anti-6×His tag antibody (Abcam, catalog # ab1187) for scFv-HER2 variants, both in 1% PBST, was added to each well for 1 h. Plates were washed three times and then developed using 50 μL 1-Step Ultra TMB-ELISA substrate solution (ThermoFisher).

### RNase A activity assay

The enzymatic activity of RNase A variants was assayed using RNaseAlert^®^-1 Kit (Integrated DNA Technologies) according to the manufacturer’s protocol. Each of the 80-times-diluted supernatant samples were normalized to have an OD_600_ equivalent to the positive control strain expressing wt RNase A. Samples were then mixed with 20 pmol of RNase A substrate and 10 μL of 10× RNaseAlert Buffer and incubated in RNase-free black 96 well microplates (Fisher) at 37 °C for 30 min. Fluorescence values were measured at 490 nm/520 nm excitation/emission wavelengths. **Thermal stability analysis.** Far-UV CD spectroscopy of purified Im7 (50-mM sodium phosphate, 400-mM sodium sulfate, pH 7.4) as a function of temperature was carried out in a 0.1-cm cuvette on a spectropolarimeter. Far-UV CD spectra were acquired between 200 nm and 260 nm with a step resolution of 1 nm. Melting temperatures of purified glycovariants was determined using high-throughput DSF as previously described (63). Briefly, 5–10 μg of proteins were mixed with Protein Thermal Shift^™^ Buffer and Protein Thermal Shift^™^ Dye purchased as Protein Thermal Shift Dye Kit^™^ (Thermo Fischer Scientific) according to manufacturer’s instructions. A melting curve was generated by monitoring fluorescence at 465 nm/610 nm excitation/emission wavelengths while increasing temperature from 10 °C to 90 °C at a rate of 0.06 °C/s on an Applied Biosystem ViiA 7 instrument (Life Technologies). To calculate *T*_m_ values, the collected data were analyzed by nonlinear regression analysis using the Boltzmann equation in Prism 8.4.2 (GraphPad).

### Computational analyses

For all computational analyses including protein structure preparation, geometric calculations, and Rosetta protocols, see **Supplementary Methods**.

## Supporting information

Supplementary Results, Methods and Figures

## Acknowledgements

We thank Markus Aebi for providing strain CLM24 and hR6 serum used in this work. We thank the Biotechnology Resource Center Genomics and Bioinformatics Core Facilities at the Cornell Institute of Biotechnology for help with sequencing experiments. The authors also thank Mike Jewett, Milan Mrksich, Eric Sundberg, Sophia Hulbert, José-Marc Techner, Weston Kightlinger, Liang Lin, Jessica Stark, and Sai Pooja Mahajan for helpful discussions of the manuscript. This work was supported by the Defense Threat Reduction Agency (HDTRA1-15-10052 and HDTRA1-20-10004 to M.P.D.), National Science Foundation (CBET-1159581, CBET-1264701, CBET-1936823 to M.P.D.), and National Institutes of Health (1R01GM137314 to M.P.D., 1R01GM127578 to M.P.D. and J.J.G., and 1S10 OD017992-01 to S.Z.). The work was also supported by seed project funding (to M.P.D.) through the National Institutes of Health-funded Cornell Center on the Physics of Cancer Metabolism (supporting grant 1U54CA210184). T.D.M was supported by a training grant from the National Institutes of Health NIBIB (T32EB023860). The content is solely the responsibility of the authors and does not necessarily represent the official views of the National Cancer Institute or the National Institutes of Health. T.J. was supported by a Royal Thai Government Fellowship and also a Cornell Fleming Graduate Scholarship.

## Author Contributions

M.L. and X.Z. designed and performed all research, analyzed all data, and wrote the paper. S.S., T.J., T.D.M., S.W.H., I.K., J.B., E.C.C., J.W.L., and J.J.G. designed and performed research, and analyzed data. Q.F. and S.Z. performed MS analysis and analyzed MS data. M.P.D. directed research, analyzed data, and wrote the paper.

## Data availability

All data generated or analyzed during this study are included in this article (and its supplementary information) or are available from the corresponding authors on reasonable request.

## Competing Interests

M.P.D. has a financial interest in Glycobia, Inc., SwiftScale Biologics, Inc., and Versatope Therapeutics, Inc. M.P.D.’s interests are reviewed and managed by Cornell University in accordance with their conflict of interest policies. J.J.G. is an unpaid board member of the Rosetta Commons. Under institutional participation agreements between the University of Washington, acting on behalf of the Rosetta Commons, Johns Hopkins University may be entitled to a portion of revenue received on licensing Rosetta software including some methods developed in this article. As a member of the Scientific Advisory Board, J.J.G. has a financial interest in Cyrus Biotechnology. Cyrus Biotechnology distributes the Rosetta software, which may include methods mentioned in this article. J.J.G.’s arrangements have been reviewed and approved by the Johns Hopkins University in accordance with its conflict of interest policies. All other authors declare no competing interests.

## References

1. G. A. Khoury, R. C. Baliban, C. A. Floudas, Proteome-wide post-translational modification statistics: frequency analysis and curation of the swiss-prot database. Sci Rep 1 (2011).

2. C. T. Walsh, S. Garneau-Tsodikova, G. J. Gatto, Jr., Protein posttranslational modifications: the chemistry of proteome diversifications. Angew Chem Int Ed Engl 44, 7342–7372 (2005).

3. M. Abu-Qarn, J. Eichler, N. Sharon, Not just for Eukarya anymore: protein glycosylation in Bacteria and Archaea. Curr Opin Struct Biol 18, 544–550 (2008).

4. A. Varki, Biological roles of glycans. Glycobiology 27, 3–49 (2017).

5. N. Mitra, S. Sinha, T. N. Ramya, A. Surolia, N-linked oligosaccharides as outfitters for glycoprotein folding, form and function. Trends Biochem Sci 31, 156–163 (2006).

6. F. S. van de Bovenkamp et al., Adaptive antibody diversification through N-linked glycosylation of the immunoglobulin variable region. Proc Natl Acad Sci U S A 115, 1901–1906 (2018).

7. G. T. Beckham et al., Harnessing glycosylation to improve cellulase activity. Curr Opin Biotechnol 23, 338–345 (2012).

8. L. X. Wang, X. Tong, C. Li, J. P. Giddens, T. Li, Glycoengineering of antibodies for modulating functions. Annu Rev Biochem 88, 433–459 (2019).

9. F. Berti, R. Adamo, Antimicrobial glycoconjugate vaccines: an overview of classic and modern approaches for protein modification. Chem Soc Rev 47, 9015–9025 (2018).

10. L. Van Landuyt, C. Lonigro, L. Meuris, N. Callewaert, Customized protein glycosylation to improve biopharmaceutical function and targeting. Curr Opin Biotechnol 60, 17–28 (2019).

11. C. J. Bosques, S. M. Tschampel, R. J. Woods, B. Imperiali, Effects of glycosylation on peptide conformation: a synergistic experimental and computational study. J Am Chem Soc 126, 8421–8425 (2004).

12. D. Shental-Bechor, Y. Levy, Effect of glycosylation on protein folding: a close look at thermodynamic stabilization. Proc Natl Acad Sci U S A 105, 8256–8261 (2008).

13. J. R. Rich, S. G. Withers, Emerging methods for the production of homogeneous human glycoproteins. Nat Chem Biol 5, 206–215 (2009).

14. T. V. Flintegaard et al., N-glycosylation increases the circulatory half-life of human growth hormone. Endocrinology 151, 5326–5336 (2010).

15. N. Ceaglio, M. Etcheverrigaray, R. Kratje, M. Oggero, Novel long-lasting interferon alpha derivatives designed by glycoengineering. Biochimie 90, 437–449 (2008).

16. A. Lusch et al., Development and analysis of alpha 1-antitrypsin neoglycoproteins: the impact of additional N-glycosylation sites on serum half-life. Mol Pharm 10, 2616–2629 (2013).

17. R. Song, D. A. Oren, D. Franco, M. S. Seaman, D. D. Ho, Strategic addition of an N-linked glycan to a monoclonal antibody improves its HIV-1-neutralizing activity. Nat Biotechnol 31, 1047–1052 (2013).

18. T. Buskas, S. Ingale, G. J. Boons, Glycopeptides as versatile tools for glycobiology. Glycobiology 16, 113R–136R (2006).

19. R. J. Payne, C. H. Wong, Advances in chemical ligation strategies for the synthesis of glycopeptides and glycoproteins. Chem Commun (Camb) 46, 21–43 (2010).

20. M. M. Chen et al., Perturbing the folding energy landscape of the bacterial immunity protein Im7 by site-specific N-linked glycosylation. Proc Natl Acad Sci U S A 107, 22528–22533 (2010).

21. A. A. Ollis, S. Zhang, A. C. Fisher, M. P. DeLisa, Engineered oligosaccharyltransferases with greatly relaxed acceptor-site specificity. Nat Chem Biol 10, 816–822 (2014).

22. G. Zhang, S. Brokx, J. H. Weiner, Extracellular accumulation of recombinant proteins fused to the carrier protein YebF in Escherichia coli. Nat Biotechnol 24, 100–104 (2006).

23. T. J. Mansell, C. Guarino, M. P. DeLisa, Engineered genetic selection links in vivo protein folding and stability with asparagine-linked glycosylation. Biotechnology journal 8, 1445–1451 (2013).

24. C. A. Dennis et al., A structural comparison of the colicin immunity proteins Im7 and Im9 gives new insights into the molecular determinants of immunity-protein specificity. Biochem J 333 (Pt 1), 183–191 (1998).

25. T. P. Ko, C. C. Liao, W. Y. Ku, K. F. Chak, H. S. Yuan, The crystal structure of the DNase domain of colicin E7 in complex with its inhibitor Im7 protein. Structure 7, 91–102 (1999).

26. A. C. Fisher et al., Production of secretory and extracellular N-linked glycoproteins in Escherichia coli. Appl Environ Microbiol 77, 871–881 (2011).

27. L. E. Yates et al., Glyco-recoded Escherichia coli: Recombineering-based genome editing of native polysaccharide biosynthesis gene clusters. Metab Eng 53, 59–68 (2019).

28. M. Wacker et al., N-linked glycosylation in Campylobacter jejuni and its functional transfer into E. coli. Science 298, 1790–1793 (2002).

29. F. Schwarz et al., A combined method for producing homogeneous glycoproteins with eukaryotic N-glycosylation. Nat Chem Biol 6, 264–266 (2010).

30. M. Kanwar, R. C. Wright, A. Date, J. Tullman, M. Ostermeier, Protein switch engineering by domain insertion. Methods Enzymol 523, 369–388 (2013).

31. A. J. Petrescu, A. L. Milac, S. M. Petrescu, R. A. Dwek, M. R. Wormald, Statistical analysis of the protein environment of N-glycosylation sites: implications for occupancy, structure, and folding. Glycobiology 14, 103–114 (2004).

32. J. D. Valderrama-Rincon et al., An engineered eukaryotic protein glycosylation pathway in Escherichia coli. Nat Chem Biol 8, 434–436 (2012).

33. M. Kowarik et al., N-linked glycosylation of folded proteins by the bacterial oligosaccharyltransferase. Science 314, 1148–1150 (2006).

34. J. M. Silverman, B. Imperiali, Bacterial N-glycosylation efficiency Is dependent on the structural context of target sequons. J Biol Chem 291, 22001–22010 (2016).

35. S. M. Juraja et al., Engineering of the Escherichia coli Im7 immunity protein as a loop display scaffold. Protein Eng Des Sel 19, 231–244 (2006).

36. J. J. Lavinder, S. B. Hari, B. J. Sullivan, T. J. Magliery, High-throughput thermal scanning: a general, rapid dye-binding thermal shift screen for protein engineering. J Am Chem Soc 131, 3794–3795 (2009).

37. R. L. Williams, S. M. Greene, A. McPherson, The crystal structure of ribonuclease B at 2.5-A resolution. J Biol Chem 262, 16020–16031 (1987).

38. U. Arnold, R. Ulbrich-Hofmann, Kinetic and thermodynamic thermal stabilities of ribonuclease A and ribonuclease B. Biochemistry 36, 2166–2172 (1997).

39. R. Grafl, K. Lang, H. Vogl, F. X. Schmid, The mechanism of folding of pancreatic ribonucleases is independent of the presence of covalently linked carbohydrate. J Biol Chem 262, 10624–10629 (1987).

40. R. Gupta, S. Brunak, Prediction of glycosylation across the human proteome and the correlation to protein function. Pac Symp Biocomput, 310–322 (2002).

41. M. M. Chen, K. J. Glover, B. Imperiali, From peptide to protein: comparative analysis of the substrate specificity of N-linked glycosylation in C. jejuni. Biochemistry 46, 5579–5585 (2007).

42. K. I. Panov et al., Ribonuclease A mutant His119 Asn: the role of histidine in catalysis. FEBS Lett 398, 57–60 (1996).

43. F. S. van de Bovenkamp et al., Variable domain N-linked glycans acquired during antigen-specific immune responses can contribute to immunoglobulin G antibody stability. Front Immunol 9, 740 (2018).

44. J. C. Krause et al., An insertion mutation that distorts antibody binding site architecture enhances function of a human antibody. mBio 2, e00345–00310 (2011).

45. W. Kightlinger et al., Design of glycosylation sites by rapid synthesis and analysis of glycosyltransferases. Nat Chem Biol 14, 627–635 (2018).

46. J. M. Techner et al., High-throughput synthesis and analysis of intact glycoproteins using SAMDI-MS. Anal Chem 92, 1963–1971 (2020).

47. S. B. Whittaker, G. R. Spence, J. Gunter Grossmann, S. E. Radford, G. R. Moore, NMR analysis of the conformational properties of the trapped on-pathway folding intermediate of the bacterial immunity protein Im7. J Mol Biol 366, 1001–1015 (2007).

48. C. Ruiz-Canada, D. J. Kelleher, R. Gilmore, Cotranslational and posttranslational N-glycosylation of polypeptides by distinct mammalian OST isoforms. Cell 136, 272–283 (2009).

49. G. A. Weiss, C. K. Watanabe, A. Zhong, A. Goddard, S. S. Sidhu, Rapid mapping of protein functional epitopes by combinatorial alanine scanning. Proc Natl Acad Sci U S A 97, 8950–8954 (2000).

50. L. M. Gregoret, R. T. Sauer, Additivity of mutant effects assessed by binomial mutagenesis. Proc Natl Acad Sci U S A 90, 4246–4250 (1993).

51. K. L. Morrison, G. A. Weiss, Combinatorial alanine-scanning. Curr Opin Chem Biol 5, 302–307 (2001).

52. P. M. Rudd et al., Glycoforms modify the dynamic stability and functional activity of an enzyme. Biochemistry 33, 17–22 (1994).

53. S. R. Hanson et al., The core trisaccharide of an N-linked glycoprotein intrinsically accelerates folding and enhances stability. Proc Natl Acad Sci U S A 106, 3131–3136 (2009).

54. R. L. Shields et al., Lack of fucose on human IgG1 N-linked oligosaccharide improves binding to human Fcgamma RIII and antibody-dependent cellular toxicity. J Biol Chem 277, 26733–26740 (2002).

55. W. Kightlinger et al., A cell-free biosynthesis platform for modular construction of protein glycosylation pathways. Nat Commun 10, 5404 (2019).

56. A. Natarajan et al., Engineering orthogonal human O-linked glycoprotein biosynthesis in bacteria. Nat Chem Biol 16, 1062–1070 (2020).

57. F. S. van de Bovenkamp, L. Hafkenscheid, T. Rispens, Y. Rombouts, The emerging importance of IgG Fab glycosylation in immunity. J Immunol 196, 1435–1441 (2016).

58. M. F. Feldman et al., Engineering N-linked protein glycosylation with diverse O antigen lipopolysaccharide structures in Escherichia coli. Proc Natl Acad Sci U S A 102, 3016–3021 (2005).

59. A. A. Ollis et al., Substitute sweeteners: diverse bacterial oligosaccharyltransferases with unique N-glycosylation site preferences. Sci Rep 5, 15237 (2015).

60. M. P. Robinson et al., Efficient expression of full-length antibodies in the cytoplasm of engineered bacteria. Nat Commun 6, 8072 (2015).

61. T. Kortemme et al., Computational redesign of protein-protein interaction specificity. Nat Struct Mol Biol 11, 371–379 (2004).

62. T. Jaroentomeechai et al., Single-pot glycoprotein biosynthesis using a cell-free transcription-translation system enriched with glycosylation machinery. Nat Commun 9, 2686 (2018).

63. U. B. Ericsson, B. M. Hallberg, G. T. Detitta, N. Dekker, P. Nordlund, Thermofluor-based high-throughput stability optimization of proteins for structural studies. Anal Biochem 357, 289–298 (2006).

