## Supplementary Results, Methods and Figures for "Shotgun scanning glycomutagenesis: a simple and efficient strategy for constructing and characterizing neoglycoproteins"

**Detailed computational analyses.** We performed computational analyses to examine the effect of site-specific glycosylation on stability and activity for three proteins, namely Im7, RNase A, and scFv-HER2. We used geometric measures and Rosetta-based calculations to identify factors behind the experimentally observed changes in the binding or catalytic activity for sequon-substituted and glycosylated variants. Complete data of all metrics plotted against experimental results can be found in **Supplementary Figs. 9** and **10** for aglycosylated and glycosylated variants, respectively, while decomposition of the Rosetta energy terms can be found in **Supplementary Figs. 11** and **12** for aglycosylated and glycosylated variants, respectively. Explanations of all measures can be found in the Materials and Methods.

For simplicity, we used “activity” to describe Im7 binding to bacterial toxin ColE7, RNase A cleavage of substrate RNA, and scFv-HER2 binding to HER2-ED. In the supplementary figures, $a_{\mathrm{wt}}$ was the activity of the wild-type (wt) protein, $a_i$ was the activity of the aglycosylated sequon-substituted variant having glycosylation site at residue $i$, and $a_{\mathrm{gly}}$ was the activity of the glycosylated sequon-substituted variant. To examine fold-changes, we examined the activity ratios of: aglycosylated sequon-substituted variant relative to wt ($a_i/a_{\mathrm{wt}}$); glycosylated sequon-substituted variant relative to the aglycosylated sequon-substituted variant ($a_{\mathrm{gly}}/a_{\mathrm{wt}}$); and glycosylated sequon-substituted variant relative to the aglycosylated sequon-substituted variant ($a_{\mathrm{gly}}/a_i$). These values were plotted on a log scale.

**Im7.** Im7 is an immunity protein whose function is to inhibit the bacterial toxin protein ColE7, so the activity for Im7 is defined by binding to ColE7. Secondary structure-specific analysis showed that a sequon substitution on a loop improved the binding activity for 48% of sequon positions, while for α-helices this value was 28% (**Supplementary Figs. 9a**). After glycosylation, binding activity was improved for 67% and 34% of residues in loops and α-helices, respectively (**Supplementary Figs. 10a**). That is, we found that loops were more receptive to glycosylation, consistent with previous studies on the location and protein environment of glycosylation sites (1-3) and with known preferences of PglB (4).

In Im7, 50% of the sequon substitutions in helix α4 resulted in improved binding activity, whereas this value was 23%, 25%, and 0% for helices α1, α2, and α3, respectively (**Supplementary Fig. 14a**). NMR studies showed that Im7 has multiple folding intermediates, with non-native contacts possible in alternate structures (5). Sequon substitutions may be altering these pathways, and mutations in α4 that improved packing (better attractive van der Waals scores) may alter the equilibrium toward the functional native state. We examined why sequon mutations at positions 52–57 (all in α3) lacked any binding activity. Helix α3 has a set of hydrophobic residues that interacts with the hydrophobic core of Im7 and polar residues that form the ColE7-binding site. Thus, a glycosylation sequon on the α3 helix likely disturbed the ColE7 binding directly or altered the placement of this helix in the binding site.

For both the aglycosylated and glycosylated Im7 variants, there was no correlation of the activity ratios with burial, distance, buried surface area (aglycosylated only), or Rosetta energies. The one exception was the van der Waals attraction component of the Rosetta energy (**Supplementary Fig. 11b**), which showed a weak correlation with the aglycosylated binding activity ($R^2$ = 0.17). Glycosylated Im7 variants showed a weak correlation of the activity ratio of glycosylated to wt for the distance of the glycosylation site from the active-site binding residues ($R^2$ = 0.13) (**Supplementary Fig. 10c**), but neither of these measures correlated with the activity ratio of the glycosylated variant relative to the aglycosylated variant (**Supplementary Fig. 13c**). In another view of the distance effect, we estimated the probability that a substitution improved activity as a function of distance from the sequon to the binding partner. The distribution indicated greater probability of activity when the sequon was close (2-8 Å) or far (above 12 Å) from the active site, with a minimum around 10 Å (**Supplementary Fig. 9d**).

To understand the steric effects of glycosylation, we further selected five glycosylated variants for which experimental *N*-glycan attachment caused no effect on activity ($Im7^{N46}$), reduced activity ($Im7^{N31}$), improved activity ($Im7^{N30}$), twofold improved activity ($Im7^{N49}$), and, finally, fourfold improved activity ($Im7^{N58}$) compared to aglycosylated variants. **Supplementary Fig. 14** shows ensembles of glycan conformations for each of these variants in the context of Im7 bound to ColE7. In experiments, glycosylation of $Im7^{N46}$ had no effect on activity ($a_{\mathrm{gly}}/a_i = 1.04$) (**Supplementary Fig. 14b**). Fittingly, the carbohydrate conformational ensemble (lines in figure) also displayed no interaction with ColE7 protein. This observation supports the hypotheses that the glycan and the bound protein must interact to change the binding activity. $Im7^{N31}$ showed a broad, disordered ensemble of structures because the glycan clashed with ColE7 residues (high Rosetta energies); accordingly, the glycosylated activity was about half of the aglycosylated activity ($a_{\mathrm{gly}}/a_i = 0.55$) (**Supplementary Fig. 14c**). In contrast, the variant for neighboring position 30 had increased activity ($a_{\mathrm{gly}}/a_i = 1.36$) and the ensemble of the bound glycan showed extended glycan chains and a favorable interaction with ColE7 (**Supplementary Fig. 14d**). The carbohydrate cloud for $Im7^{N49}$ showed a few carbohydrate chains (<10%) that interacted with nearby ColE7 residues, and a stronger increase of binding activity upon glycosylation ($a_{\mathrm{gly}}/a_i = 1.91$) (**Supplementary Fig. 14e**). Finally, for $Im7^{N58}$, which had the highest increase in activity upon glycosylation activity ($a_{\mathrm{gly}}/a_i = 3.44$), again, many of the low-energy glycan conformations made favorable interactions with ColE7 (**Supplementary Fig. 14f**). Thus, in these cases, the structures and interactions of the *N*-glycans on Im7 with ColE7 explains the change in binding activity upon glycosylation. For these cases, the Rosetta total score followed these observations (colored points in **Supplementary Fig. 10f**). However, one cannot make general predictions from the Rosetta energy as there were many residues with varying activities for similar Rosetta score changes (**Supplementary Fig. 13f**) and so Rosetta energy in general did not correlate with the activity ratios.

**RNase A.** All of the analysis of RNase A is given in the main Results section, as the disruption of active site residues or disulfide bonds explained most effects. Indeed, RNase A active site residues (Gln11, His12, Lys41, His119, Phe120, Asp121) and substrate specificity determining residues (Thr45, Asn71, Asp83, Glu111) are disparately located throughout the primary sequence (6, 7) as well as the four disulfide bonds (Cys26-Cys84, Cys40-Cys95, Cys58-Cys110, and Cys65-Cys72) that make important contributions to RNase A activity and structural rigidity (8).

**Anti-HER2 scFv.** ScFv-HER2 is an antibody specific for HER2, therefore the activity is defined by binding to HER2. Concerning the secondary structure, we found that sequon substitution improved, or did not affect, the activity for approximately 40% of instances for β-strands and 62% for loops (**Supplementary Fig. 9a**). After glycosylation, these numbers changed to 50% and 60%, respectively (**Supplementary Fig. 10a**), indicating that like Im7, scFv-HER2 loops are better positions for achieving improved binding through glycosylation. Like the Im7–ColE7 case, the probability of increasing activity as a function of distance from the sequon to the binding site was high in a close region (5 to 18 Å) and a far region (over 28 Å, **Supplementary Fig. 9d**). However, glycosylation reduced these regions to 10 to 19 Å and over 22 Å (**Supplementary Fig. 10d**).

Compared to the other two systems, scFv-HER2 showed correlations with more geometric and Rosetta measures. Most strikingly, the change in Rosetta score correlated with the ratio of aglycosylated activity to wt ($R^2$ = 0.49, **Supplementary Fig. 9f**), and the ratio of glycosylated activity to wt ($R^2$ = 0.49, **Supplementary Fig. 10f**). For aglycosylated and glycosylated activities, improved interface energy showed correlation with better binding ($R^2$ = 0.18 for aglycosylated, **Supplementary Fig. 9g,** and $R^2$ = 0.63 for glycosylated, **Supplementary Fig. 10g**). The individual Rosetta score terms that correlated with the activity ratios were repulsive and attractive van der Waals, and to a good extent, the electrostatics term (**Supplementary Figs. 11** and **12**).

### Supplementary Methods

**Next-generation sequencing.** SSGM libraries were prepared for sequencing using a two-step PCR strategy. First, POI gene-specific amplification was performed with 50 ng plasmid DNA and primers also containing partial Illumina adaptor sequences (forward: 5'-TCG TCG GCA GCG TCA GAT GTG TAT AAG AGA CAG NNN NCG GAA TAT CAG CGG CGT TCT AG-3', reverse: 5'-GTC TCG TGG GCT CGG AGA TGT GTA TAA GAG ACA GNN NNC CTT GTA GTC TGC TCC GTC G-3') using the following thermocycler conditions: 95 °C for

3 min, (95 °C for 30 s, 53 °C for 30 s, 72 °C for 30 s) x 25 cycles, 72 °C for 5 min. Different forward and reverse primers were used to amplify the HER2 heavy and light variable regions (forward: 5'-TCG TCG GCA GCG TCA GAT GTG TAT AAG AGA CAG NNN NGG CGG TGG CGG ATC G -3', reverse: 5'-GTC TCG TGG GCT CGG AGA TGT GTA TAA GAG ACA GNN NNC CTG AAC CGC CTC CAC C-3'). A second PCR was performed to add full-length Nextera adaptors using 20 ng of the first reaction using the following thermocycler conditions: 95 °C for 2 min, (95 °C for 30 s, 50 °C for 30 s, 72 °C for 30 s) x 8 cycles, 72 °C for 5 min. Amplicons were quantified using a Qubit 3.0 Fluorimeter (ThermoFisher), pooled, and submitted for paired-end 2 x 250 bp sequencing on the MiSeq instrument (Illumina). The resulting reads were trimmed and merged using PANDAseq followed by collapsing of identical sequences using FASTX. A custom Python script, available upon request, was used for identification of glycosites within each gene sequence.

**Mass spectrometry analysis of protein glycosylation.** Proteins samples (~2 μg) were separated by SDS-PAGE gel and bands corresponding to glycosylated YebF-Im7$^{DQNAT}$ were excised and subjected to in-gel digestion by trypsin followed by extraction of tryptic peptides essentially as described (9). Gel slices were washed and then destained by treatment with a 1:1 mixture of acetonitrile (Fisher Chemical) and 50 mM aqueous $NH_4HCO_3$ followed by treatment with 100% acetonitrile. After destaining, gel pieces were reduced and alkylated with 10 mM dithiothreitol (Roche) and 50 mM iodoacetamide (Acros Organics). The glycoproteins were directly digested by adding trypsin (w/w = 1:10) in 50 mM $NH_4HCO_3$ to the gel pieces and incubating overnight at 37 °C. The tryptic peptides were extracted by 50% acetonitrile with 5% formic acid (Fisher Chemical) and 75% acetonitrile with 5% formic acid. Extracted peptides were pooled and dried by SpeedVac. The tryptic peptides were suspended in 24 μL of 0.5% formic acid and 10 μL was injected into an UltiMate3000 RSLCnano (Dionex) coupled to an Orbitrap Fusion mass spectrometer (Thermo-Fisher Scientific) as described (10) with slight modifications. The peptides were injected onto a PepMap C18 RP nano trapping column (5 μm, 100 μm i.d x 20 mm) at 20 μL/min flow rate for rapid sample loading, and separated on an Acclaim PepMap C18 nano column (3 μm, 75 μm x 25cm, Thermo Fisher Scientific). The tryptic peptides were eluted in a 90 min gradient of 5% to 23% to 35% solvent B (95% acetonitrile, 0.1% formic acid) corresponding to 3 to 73 to 93 min, respectively, at 300 nL/min. The 90-

min gradient was followed by a 9-min ramping to 90% B, a 9-min hold at 90% B and quick switch to 5% B and 95% solvent A (2% acetonitrile, 0.1% formic acid) for 1 min. The column was re-equilibrated with 95% A for 25 min prior to the next run. The Orbitrap Fusion was operated in positive ion mode with nanospray voltage set at 1.7 kV and source temperature at 275 °C. The MS survey scan was acquired at a resolving power of 120,000 (FWHM at *m*/*z* 200) across *m*/*z* 350-1800, which was followed by a "top speed" data-dependent electron-transfer dissociation (ETD) MS/MS scan (cycle time of 4 s) supplemented with higher-energy collision dissociation (EThcD) fragmentation workflow for precursor peptides with 3–6 charges. EThcD fragmentation was acquired using calibrated charge-dependent ETD parameters supplemented with 15% collisional energy in ion trap detector with automatic gain control (AGC) of $3x10^4$ and maximum injection time of 118 s.

Data analysis was performed by Byonic v3.6 (Protein Metrics) searching software against an *E. coli* database containing the YebF-Im7 protein, and an in-house generated *N*-linked glycan database with additional diBacNAc-containing glycans. The peptide search parameters were as follows: two missed cleavage for full trypsin digestion with fixed carbamidomethyl modification of cysteine and Q to Pyro-E on N-terminal Q, variable modifications of methionine oxidation and deamidation on asparagine/glutamine residues. The peptide mass tolerance was 10 ppm and fragment mass tolerance values for EThcD spectra was 0.6 Da. Both the maximum number of common and rare modifications were set at two. Identified peptides and glycopeptides were filtered for a [log Prob] value >3 and glycopeptide mass error <5 ppm. The search results were exported to excel files for final analysis and graphical presentation.

**Protein structure preparation.** Initial coordinates of structures used for analysis were obtained from the Protein Data Bank at RCSB.org as follows: the Im7–E7 complex (PDB: 2JBG), RNAse A bound to four-nucleotide-long DNA (PDB: 1RCN, DNA: ATAA) and the scFv-Her2–ErbB2 complex (PDB: 3WSQ). For RNase A, we modeled an eight-nucleotide-long RNA at the same binding site of the DNA. We used X3DNA (11) to identify nucleotide arrangement parameters from DNA, as a template, to model RNA. We replaced the thymine residue of bound DNA with its RNA counterpart uracil. Since an initial analysis showed that four-nucleotide-long RNA is small for an interface-interaction calculation, we added two extra nucleotides to both ends of the modeled RNA to make it eight nucleotides long.

Finally, to remove any unwanted clashes coming from the base sugar (deoxyribose to ribose) and the nucleotide change (thymine to uracil), we used the Rosetta relax protocol with a restraint on protein backbone atoms to generate a pool of 100 RNA–protein structures ("decoys"). By manual inspection, we removed decoys showing RNAs outside of the binding site of RNase A. From the pool of remaining structures, we selected a candidate with the best Rosetta score for our study.

**Geometric calculations.** Secondary structure was identified using the DSSP protocol in PyRosetta (12). Burial of a glycosylated residue was represented by a count of $C_\beta$ atoms within 8.5 Å from the $C_\beta$ atom of the selected amino acid residue ($C_\alpha$ for alanine). The distance of a glycosylated residue from its binding partner was estimated by the $C_\alpha$–$C_\alpha$ distance from the glycosylated residue to the closest residue of the binding partner. For RNase A, the distance of glycosylated residue from the active site residue His-119 was used instead, as RNase A does not bind to a protein. This distance was used for both the "activity ratio" and the "activity improvement probability" analyses. Interface solvent-accessible surface area (SASA) was measured using the Rosetta interface analyzer (13).

**Rosetta measures.** Rosetta's REF15 score function (14) was used to estimate the stability of a protein–protein complex. An upper-bound cutoff of 100 REU is used to limit the effect of bad conformers on the correlation, while for Im7, this value was 60 REU. To represent the binding energy of a complex, the interface score was calculated with the Rosetta Interface Analyzer (13) by subtracting the score of the separated monomers from the complex.

**Rosetta protocols for sequon substitution and glycomutagenesis.** We used the RosettaCarbohydrate framework (15) and a new glycomutagenesis protocol to input a wt PDB file and generate all possible sequon-substituted variants and glycosylated variants. Coordinate files after sequon substitution and side chain optimization steps were used as aglycosylated variants for analysis. Calculations were carried out in Rosetta release version 2020.06 and PyRosetta4 release 247 (www.rosettacommons.org). The command line for the glycomutagenesis calculation was:

```
glycomutagenesis.linuxgccrelease -in:file:s <PDB> -include_sugars
-nstruct <length of sequence selected for glycomutagenesis>
-n_cycles 100 -out:path:pdb ./output -out:path:score ./output
```

The command line for the relaxation runs was:

```
relax.linuxgccrelease -s <PDB> -nstruct 100
-relax:default_repeats 5
-relax:bb_move false
-out:path:pdb ./output_relax -out:path:score ./output_relax
```

The command for analyzing aglycosylated structures was:

```
InterfaceAnalyzer.default.linuxgccrelease -s <PDB>
-interface A_B #(For chain A and B)
-out:path:pdb ./output -out:path:score ./output
```

and the command for glycosylated structures was:

```
InterfaceAnalyzer.default.linuxgccrelease -s <pdb file>
-interface AB_C #(For [protein A + Glycan B] and Protein C)
-fixedchains A B #(For Protein A + Glycan B)
-out:path:pdb ./output -out:path:score ./output
```

**Supplementary References**


1. D. F. Zielinska, F. Gnad, J. R. Wisniewski, M. Mann, Precision mapping of an *in vivo* *N*-glycoproteome reveals rigid topological and sequence constraints. *Cell* **141**, 897-907 (2010).
2. D. F. Zielinska, F. Gnad, K. Schropp, J. R. Wisniewski, M. Mann, Mapping *N*-glycosylation sites across seven evolutionarily distant species reveals a divergent substrate proteome despite a common core machinery. *Mol Cell* **46**, 542-548 (2012).
3. A. J. Petrescu, A. L. Milac, S. M. Petrescu, R. A. Dwek, M. R. Wormald, Statistical analysis of the protein environment of *N*-glycosylation sites: implications for occupancy, structure, and folding. *Glycobiology* **14**, 103-114 (2004).
4. M. Kowarik *et al.*, *N*-linked glycosylation of folded proteins by the bacterial oligosaccharyltransferase. *Science* **314**, 1148-1150 (2006).
5. S. B. Whittaker, G. R. Spence, J. Gunter Grossmann, S. E. Radford, G. R. Moore, NMR analysis of the conformational properties of the trapped on-pathway folding intermediate of the bacterial immunity protein Im7. *J Mol Biol* **366**, 1001-1015 (2007).
6. R. T. Raines, "Active Site of Ribonuclease A" in Nucleic Acids and Molecular Biology*,* M. A. Zenkova, Ed. (Springer-Verlag Berlin Heidelberg, 2004), vol. 13, pp. 19-32.
7. S. B. DelCardayre (1994) Catalysis by Ribonuclease A: Specificity, Processivity, and Mechanism. in *Department of Biochemistry* (University of Wisconsin, Madison).
8. T. A. Klink, K. J. Woycechowsky, K. M. Taylor, R. T. Raines, Contribution of disulfide bonds to the conformational stability and catalytic activity of ribonuclease A. *Eur J Biochem* **267**, 566-572 (2000).
9. A. Natarajan *et al.*, Engineering orthogonal human *O*-linked glycoprotein biosynthesis in bacteria. *Nat Chem Biol* **16**, 1062-1070 (2020).
10. M. Song *et al.*, IRE1alpha-XBP1 controls T cell function in ovarian cancer by regulating mitochondrial activity. *Nature* **562**, 423-428 (2018).

11. X. J. Lu, W. K. Olson, 3DNA: a software package for the analysis, rebuilding and visualization of three-dimensional nucleic acid structures. *Nucleic Acids Res* **31**, 5108-5121 (2003).
12. S. Chaudhury, S. Lyskov, J. J. Gray, PyRosetta: a script-based interface for implementing molecular modeling algorithms using Rosetta. *Bioinformatics* **26**, 689-691 (2010).
13. P. B. Stranges, B. Kuhlman, A comparison of successful and failed protein interface designs highlights the challenges of designing buried hydrogen bonds. *Protein Sci* **22**, 74-82 (2013).
14. R. F. Alford *et al.*, The Rosetta all-atom energy function for macromolecular modeling and design. *J Chem Theory Comput* **13**, 3031-3048 (2017).
15. J. W. Labonte, J. Adolf-Bryfogle, W. R. Schief, J. J. Gray, Residue-centric modeling and design of saccharide and glycoconjugate structures. *J Comput Chem* **38**, 276-287 (2017).

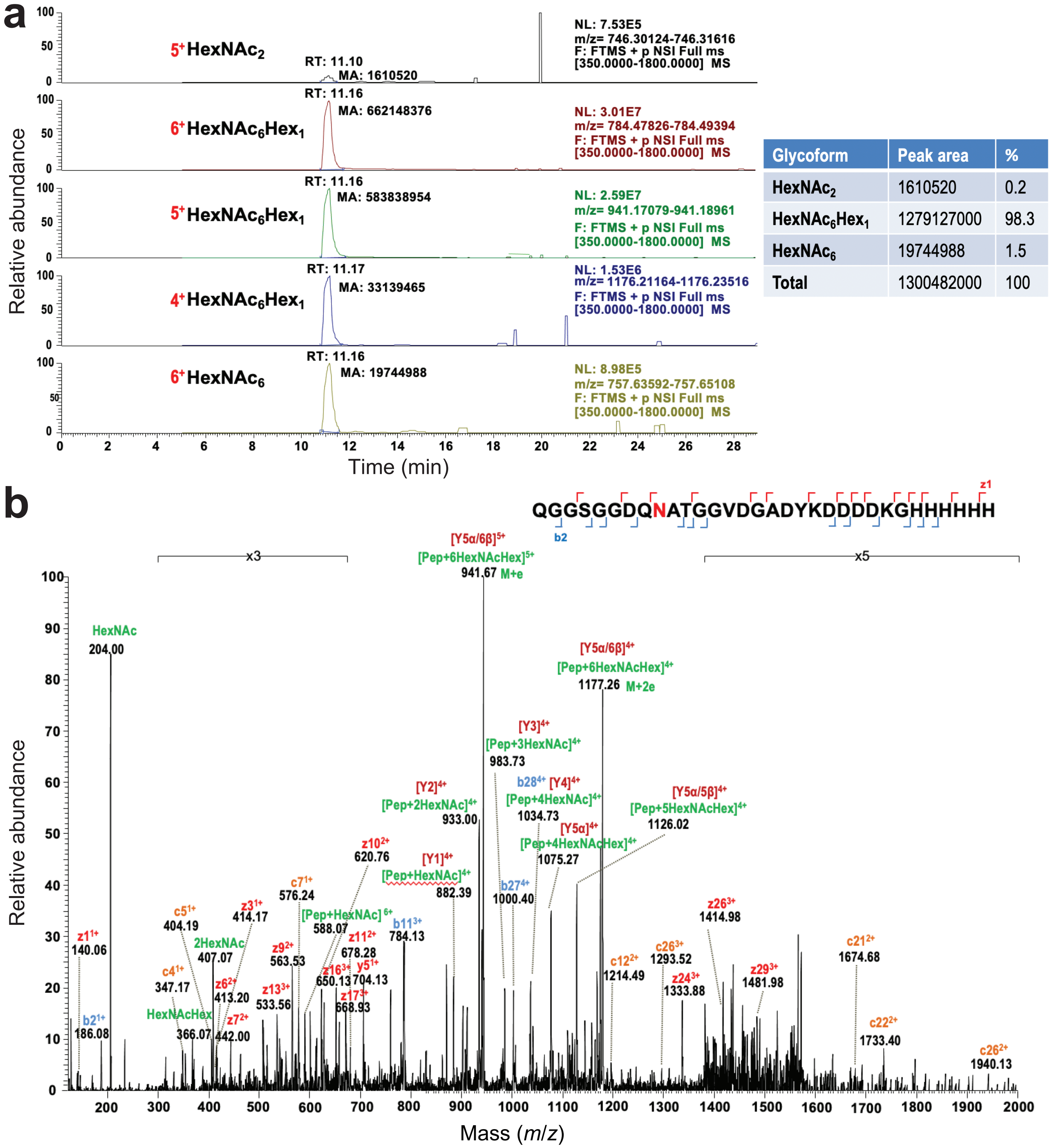


| Glycoform | Peak area | % |
|---|---|---|
| $HexNAc_2$ | 1610520 | 0.2 |
| $HexNAc_6Hex_1$ | 1279127000 | 98.3 |
| $HexNAc_6$ | 19744988 | 1.5 |
| Total | 1300482000 | 100 |

**Supplementary Figure 1. MS analysis of YebF-Im7$^{DQNAT}$ glycosylation.** (a) Extracted ion chromatograms (XIC) of three glycoforms confidently identified in the sample including $HexNAc_2$ with charge (*z*) of $5^+$ (top panel) $HexNAc_6Hex_1$ with 3 *z* ions of $6^+$, $5^+$, $4^+$ (three middle panels) and $HexNAc_6$ with *z* of $6^+$ (bottom panel) on the same core peptide QGGSGGDQNATGGVDGADYKDDDDKGHHHHHH of protein YebF-Im7$^{DQNAT}$. All three glycoforms eluted at 11.1 min with $HexNAc_6Hex_1$ serving as the predominant glycoform with an estimated abundance of >98% of the total glycoforms in the sample (table at right). (b) A representative MS/MS spectrum of a $6^+$ charged precursor ion at *m/z* 784.31 identifying the glycopeptide with $HexNAc_6Hex_1$ attached to the C-terminal DQNAT motif in YebF-Im7$^{DQNAT}$. The EThcD MS/MS spectrum reveals the desired $GalNAc_5$(Glc)GalNAc structure on the core peptide as shown in y,z and b,c series ions as well as complete Y-type series ions (from Y1 to Y5α/6β).

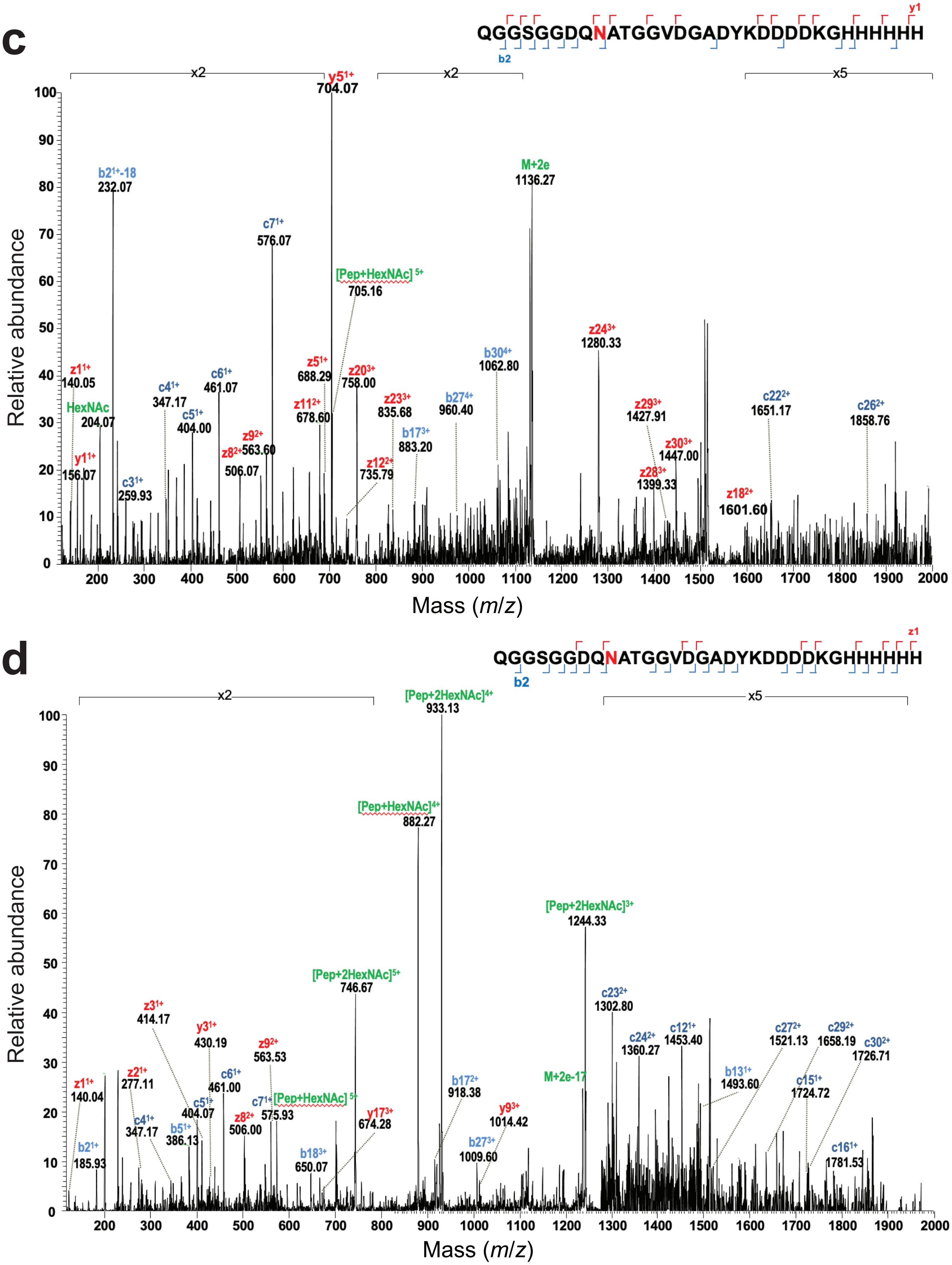


**Supplementary Figure 1. MS analysis of YebF-Im7$^{DQNAT}$ glycosylation.** Representative EThcD MS/MS spectra for two minor glycoforms identified in the sample at (c) *m/z* 757.64 with charge $6^+$ for the $HexNAc_6$ glycoform and (d) *m/z* 746.31 with charge $5^+$ for the $HexNAc_2$ glycoform with the y,z and b,c series ions similar to Supplementary Figure 1b.

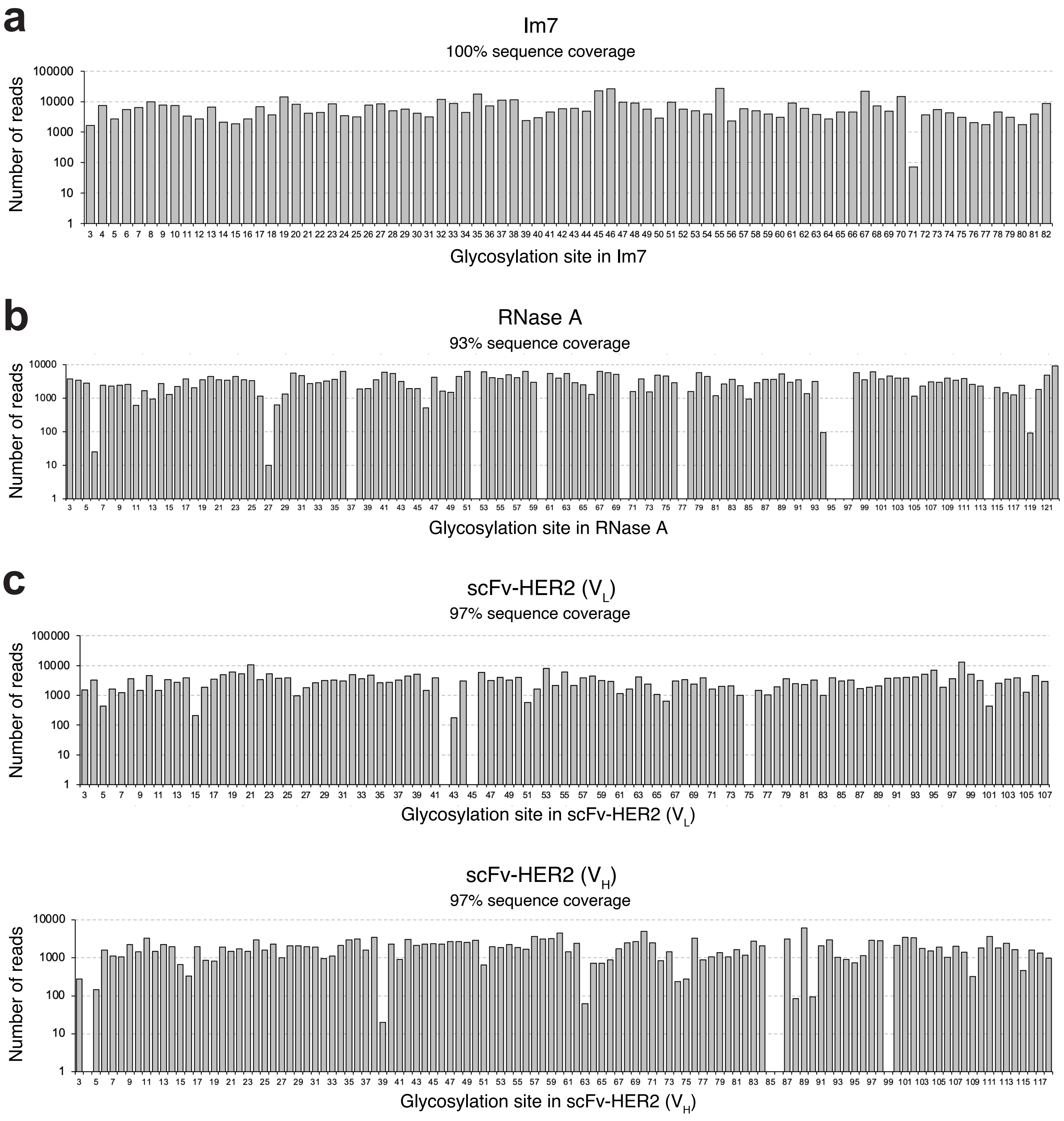


**Supplementary Figure 2. Next-generation sequencing (NGS) anslysis of SSGM libraries.** Read count of sequences containing the DQNAT sequon in (a) Im7, (b) RNAseA, and (c) scFv-HER2 libraries obtained by paired-end 2 x 250 bp MiSeq. A minimum of 800,000 reads were obtained for each library. After data processing, full-length paired sequencing reads were scanned for sequon location. Each bar represents the number of reads, at minimum 10, containing the sequon at a given location in the protein sequence, denoted by the asparagine residue position.

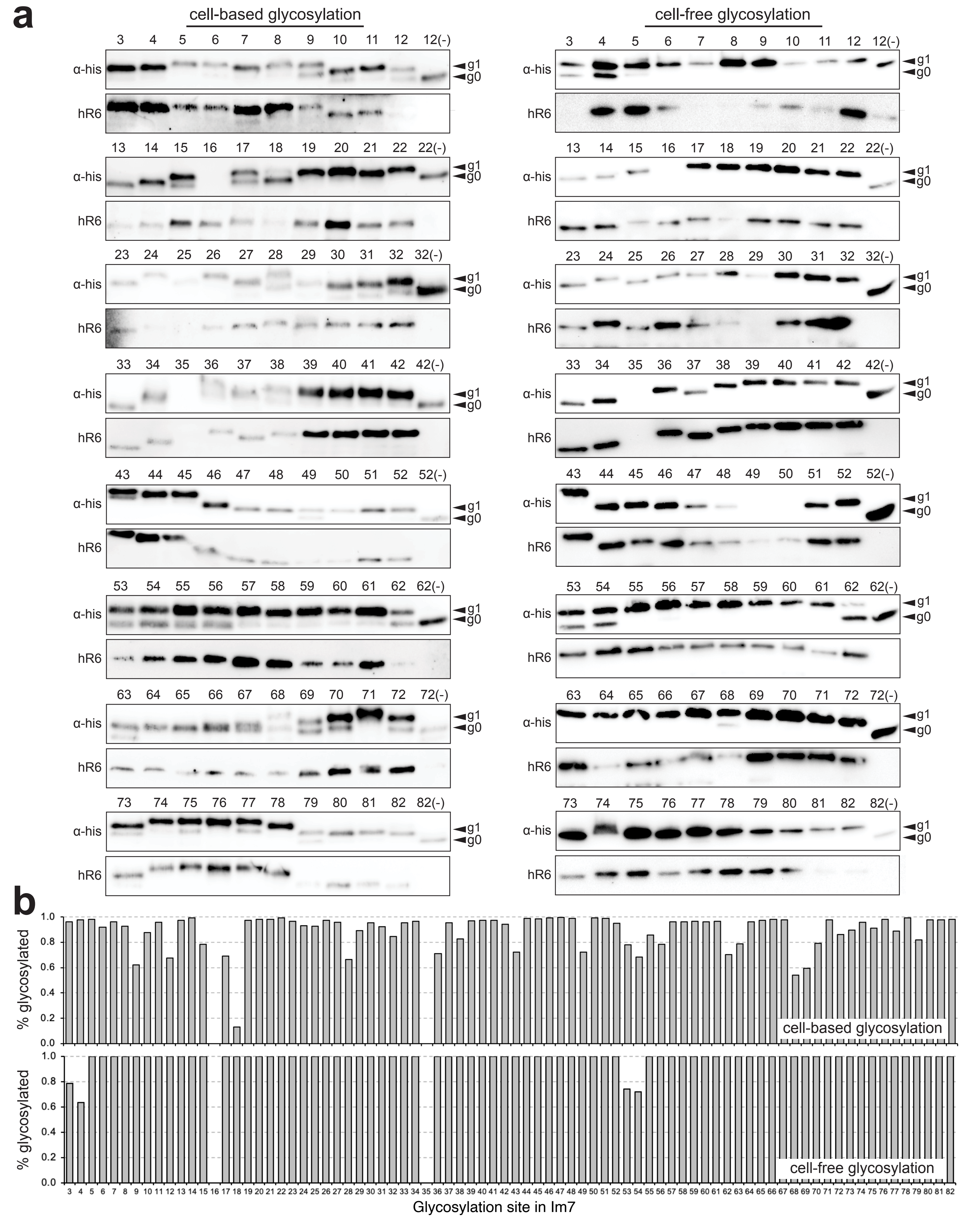


**Supplementary Figure 3. Cell-based versus cell-free glycosylation of Im7.** (a) Immunoblot analysis of cell-based (left) and cell-free (right) Im7 glycosylation. For cell-based blots, supernatant fractions were isolated from CLM24 cells carrying plasmids encoding YebF-Im7 variants (sequon mutations at the indicated position) along with requisite *N*-glycosylation machinery. For cell-free blots, aglycosylated YebF-Im7 variants were incubated with purified *Cj*PglB and extracted LLOs bearing modified *C. jejuni* glycan. Blots were probed with anti-polyhistidine antibody (α-His) to detect acceptor protein (top panel) and hR6 serum against the glycan (bottom panel). Markers for aglycosylated (g0) and singly glycosylated (g1) forms of Im7 are indicated at right. Results are representative of at least three biological replicates. (b) Glycosylation efficiency of cell-based (top panel) and cell-free (bottom panel) Im7 glycosylation where % glycosylated was calculated as the ratio g1/[g0+g1], and g0 and g1 values were determined by densitometric quantification of bands in anti-His immunoblots in (a) for each neoglycoprotein.

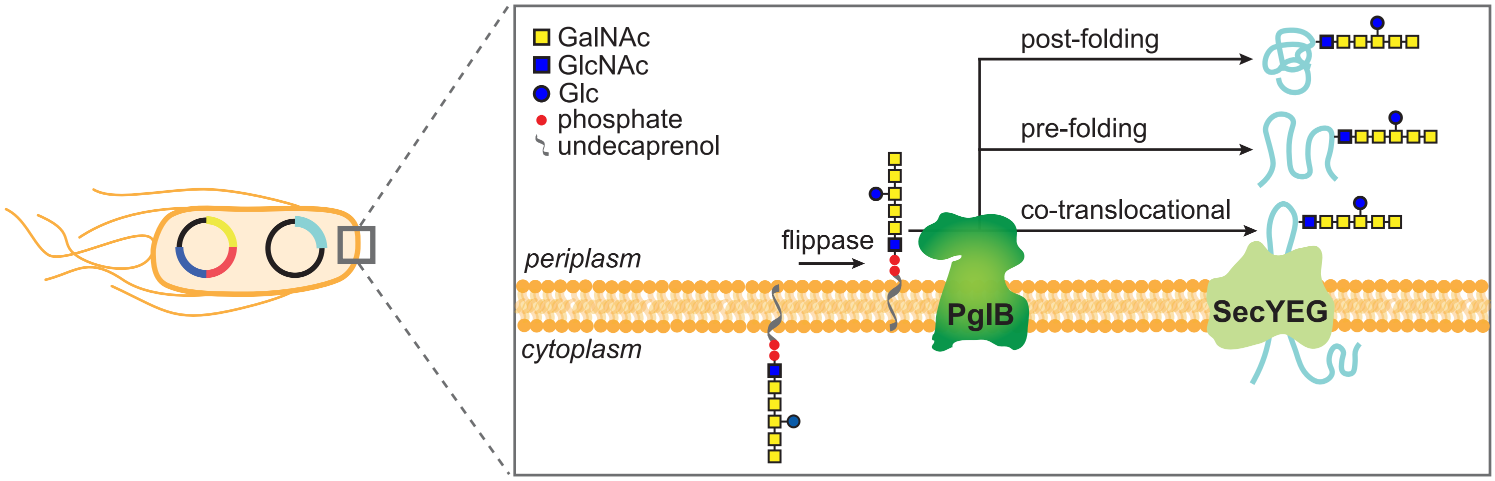


**Supplementary Figure 4. Timing of protein glycosylation in bacterial expression host.** Schematic of possible conformational states of substrate proteins during bacterial *N*-glycosylation. PglB can transfer glycans onto a substrate protein: (1) co-translocationally, before folding is complete; (2) post-translocationally, before folding is complete; and (3) post-translocationally, after folding is complete. Adapted from Silverman and Imperiali (*J Biol Chem*, 2016).

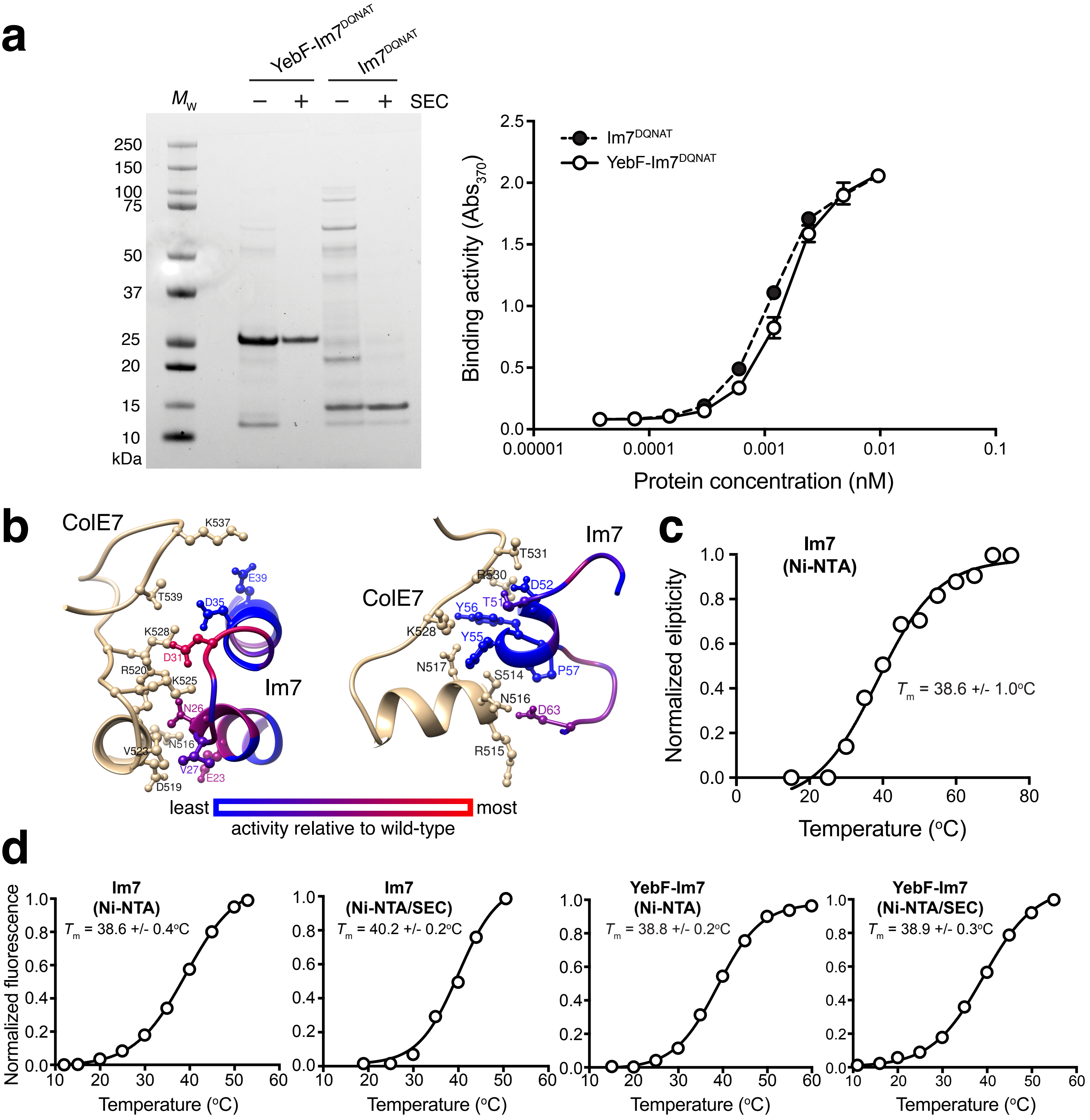


**Supplementary Figure 5. Characterization of Im7 activity and stability.** (a) Coomassie-blue stained SDS-PAGE gel of YebF-Im7$^{DQNAT}$ or unfused Im7$^{DQNAT}$ purified by Ni-NTA with (+) or without (-) additional SEC purification step. Molecular weight ($M_W$) ladder pictured at left. Binding activity of Ni-NTA-purified YebF-Im7$^{DQNAT}$ (open circle) and unfused Im7$^{DQNAT}$ (closed circle) was quantified by ELISA using ColE7 as immobilized antigen. Data are average of three biological replicates and error bars represent standard deviation of the mean. (b) Detailed interactions between ColE7 and Im7, highlighting the sidechains of Im7 in regions of α1-loop12-α2 (residue 19 to 39; middle) and loop23-α3-loop34 (residue 46 to 63; right). Heatmap analysis of change in binding activity was determined by normalizing activity measured for aglycosylated sequon variant by that of wt Im7. (c) Thermal stability of Ni-NTA-purified Im7 protein determined by CD thermal denaturation. Far-UV CD spectra were acquired between 200-260 nm with step resolution of 1 nm. (d) Thermal stability analysis of same proteins from (a) using DSF with SYPRO Orange dye in real-time PCR instrument. $T_m$ values calculated as midpoint of thermal transition between native and unfolded states. Results are representative of three biological replicates.

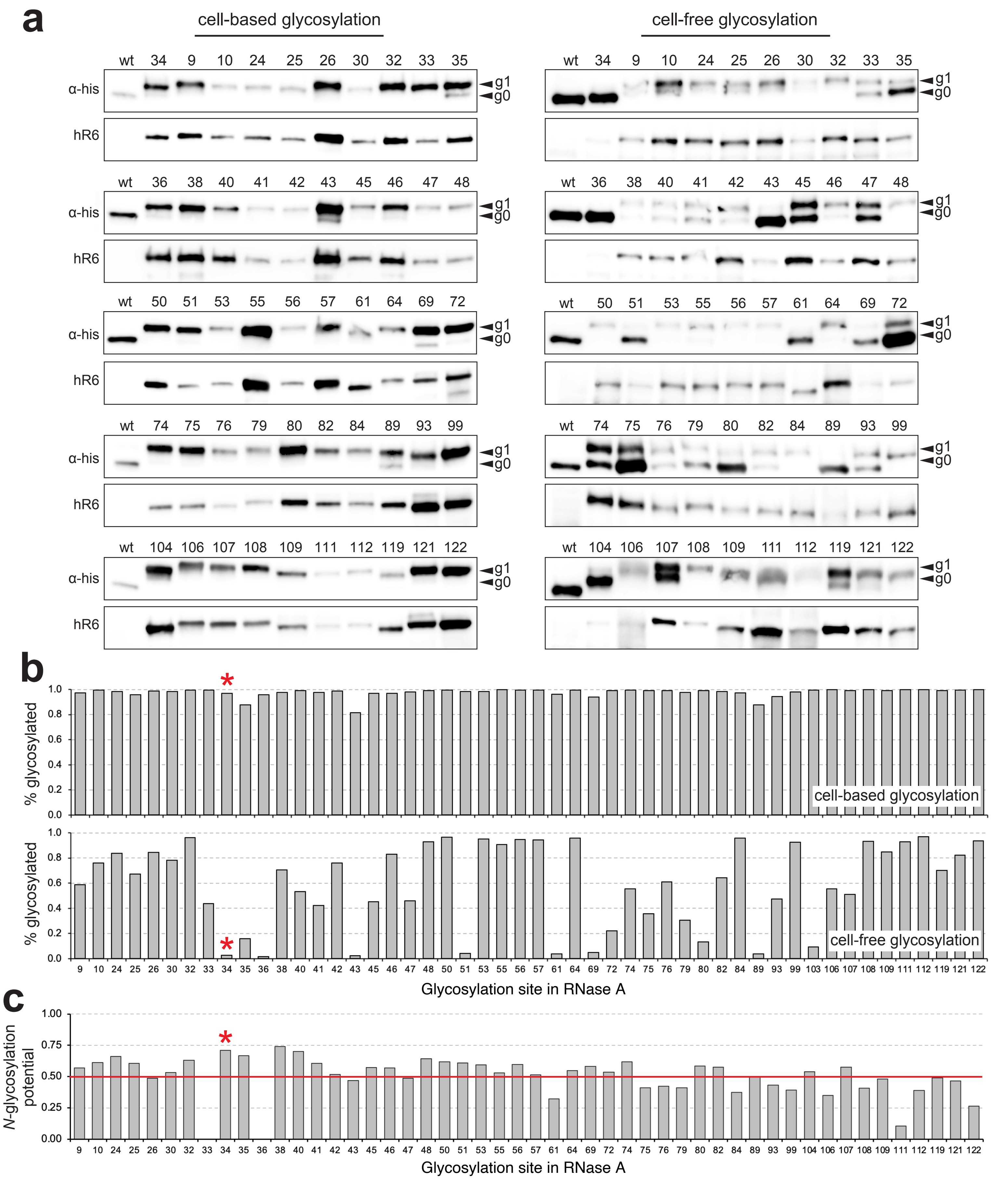


**Supplementary Figure 6. Cell-based versus cell-free glycosylation of RNase A.** Immunoblot analysis of cell-based (left) and cell-free (right) RNase A glycosylation. For cell-based blots, periplasmic fractions were isolated from CLM24 cells carrying plasmids encoding YebF-RNase A variants (sequon mutations at the indicated position) along with requisite *N*-glycosylation machinery. For cell-free blots, aglycosylated YebF-RNase A variants were incubated with purified *Cj*PglB and extracted LLOs bearing modified *C. jejuni* glycan. Blots were probed with anti-polyhistidine antibody (α-His) to detect acceptor protein (top panel) and hR6 serum against the glycan (bottom panel). Markers for aglycosylated (g0) and singly glycosylated (g1) forms of acceptor protein are indicated at right. Results are representative of at least three biological replicates. (b) Glycosylation efficiency of cell-based (top panel) and cell-free (bottom panel) RNase A glycosylation where % glycosylated was calculated as the ratio g1/[g0+g1], and g0 and g1 values were determined by densitometric quantification of bands in anti-His immunoblots in (a) for each neoglycoprotein.(c) Predicted *N*-glycosylation potential of each glycosite in RNase A using NetNGlyc1.0 server tool (http://www.cbs.dtu.dk/services/NetNGlyc/). Any potential crossing above the default threshold value of 0.5 (red line) represents a predicted glycosylated site. Red asterisk indicates native N34 glycosite.

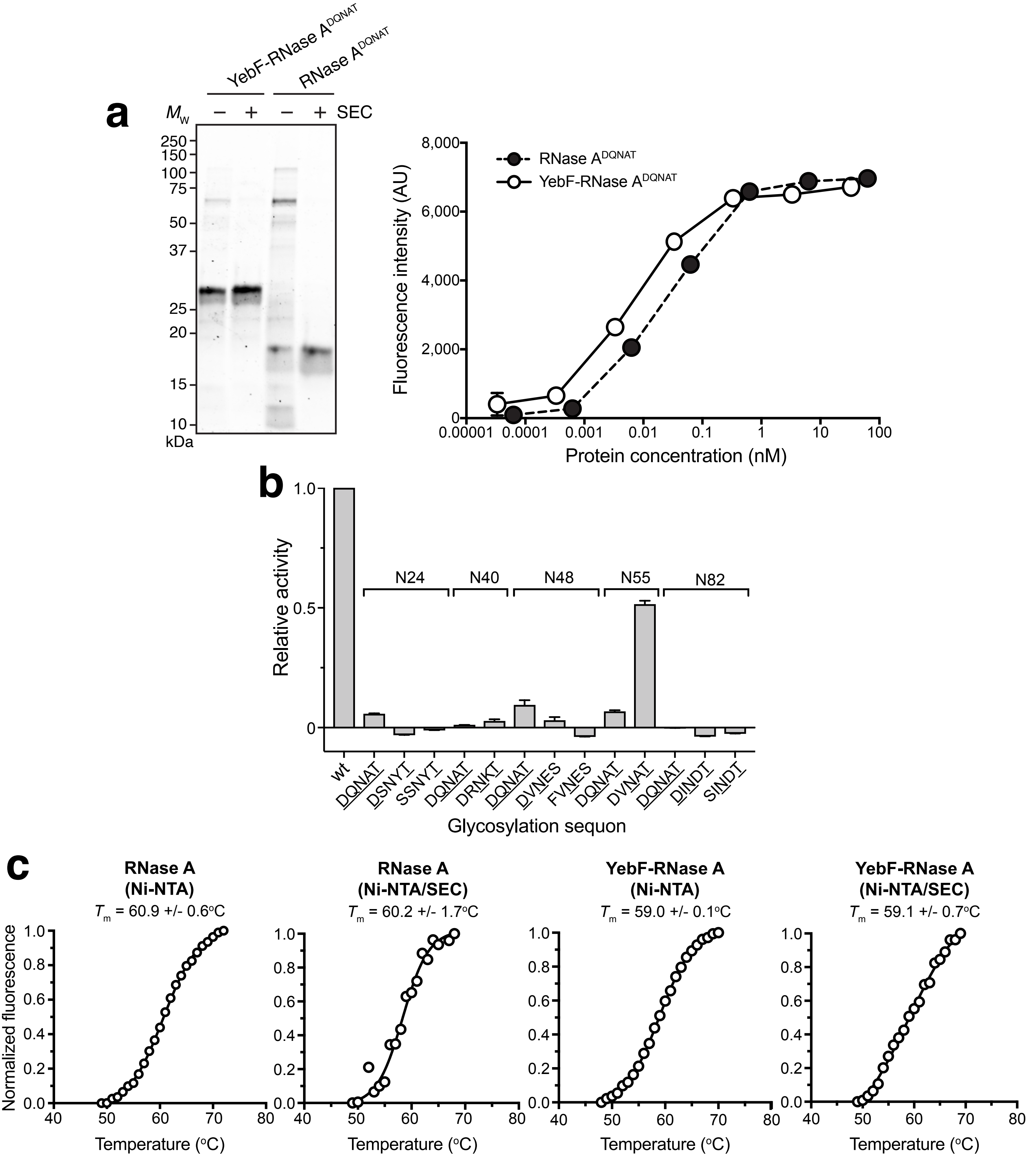


**Supplementary Figure 7. Characterization of RNase A activity and stability.** (a) Coomassie-blue stained SDS-PAGE gel of YebF-RNase $A^{DQNAT}$ or unfused RNase $A^{DQNAT}$ purified by Ni-NTA with (+) or without (-) additional SEC purification step. Molecular weight ($M_W$) ladder pictured at left. RNase activity of Ni-NTA-purified YebF-RNase $A^{DQNAT}$ (open circle) and unfused RNase $A^{DQNAT}$ (closed circle). Data are average of three biological replicates and error bars represent standard deviation of the mean. (b) RNase activity of aglycosylated versions of minimal sequon mutants that were generated by more conservative mutation of RNase A. At each indicated position (N24, N40, N48, N55, and N82), typically one or two mutations (indicated by underlined letters) were introduced by site-directed mutagenesis to yield minimal sequion, either D-X-N-X-T/S or X-X-N-X-T/S (where X represents the native amino acid). All data were normalized to the binding activity measured for aglycosylated wild-type RNase A (wt). Data are average of three biological replicates and error bars represent standard deviation of the mean. (c) Thermal stability analysis of same proteins from (a) using DSF with SYPRO Orange dye in real-time PCR instrument. $T_m$ values calculated as midpoint of thermal transition between native and unfolded states. Results are representative of three biological replicates.

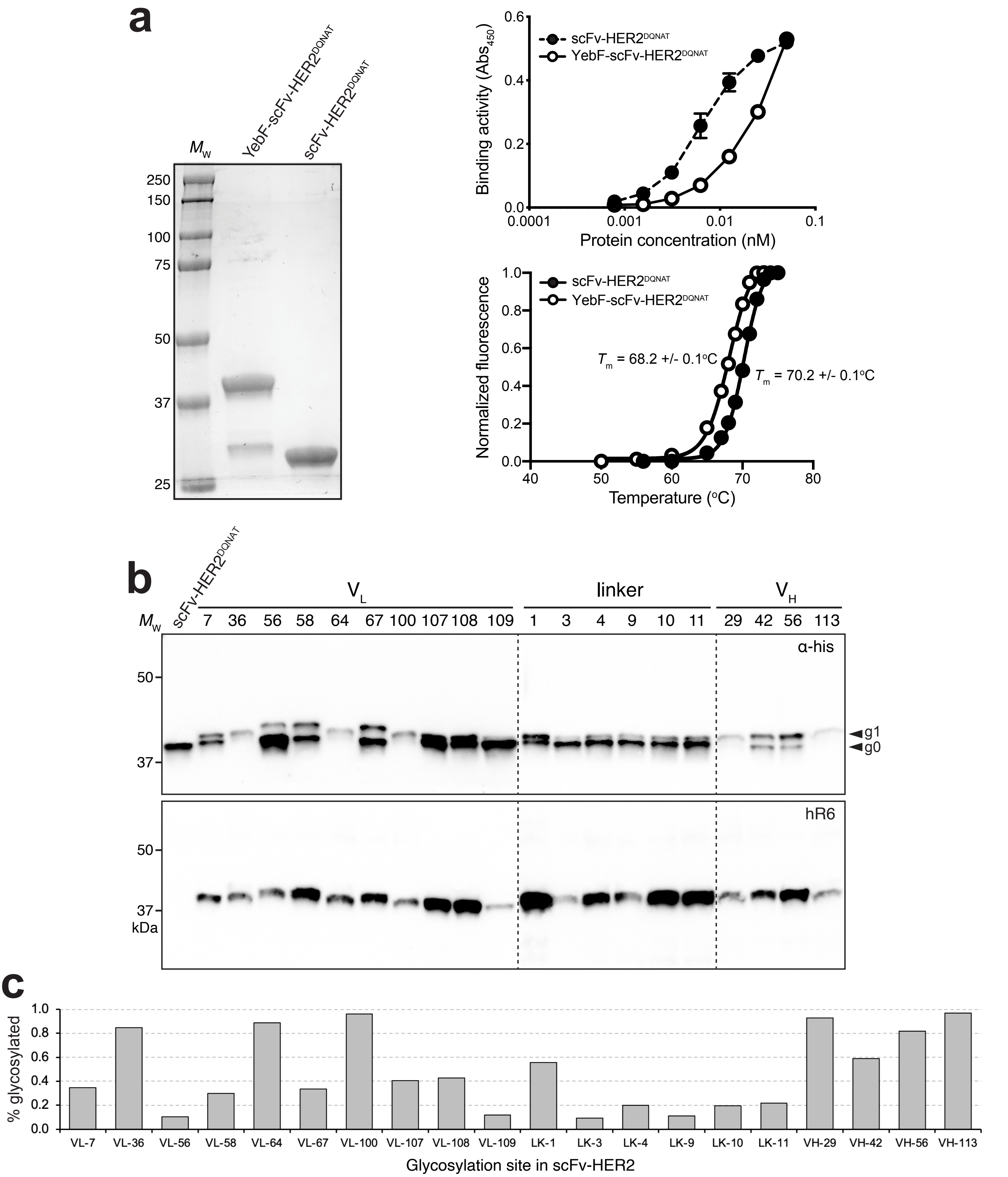


**Supplementary Figure 8. Characterization of scFv-HER2 activity and glycosylation.** (a) (left) Coomassie-blue stained SDS-PAGE gel of YebF-scFv-HER2$^{DQNAT}$ or unfused scFv-HER2$^{DQNAT}$ purified by Ni-NTA. Molecular weight ($M_W$) ladder pictured at left. (right, top) Antigen-binding activity of Ni-NTA-purified YebF-scFv-HER2$^{DQNAT}$ (open circle) and unfused scFv-HER2$^{DQNAT}$ (closed circle) was quantified by ELISA using HER2-ED as immobilized antigen. (Right, bottom) Thermal stability analysis of same proteins using DSF with SYPRO Orange dye in real-time PCR instrument. $T_m$ values calculated as midpoint of thermal transition between native and unfolded states. All results are representative of three biological replicates. (b) Western blot analysis of periplasmic fractions from CLM24 cells carrying plasmids encoding YebF-scFv-HER2 fusions with sequon mutations at the indicated position along with requisite *N*-glycosylation machinery. Blots were probed with anti-polyhistidine antibody (α-His) to detect acceptor protein (top panel) and hR6 serum against the glycan (bottom panel). Markers for aglycosylated (g0) and singly glycosylated (g1) forms of acceptor protein are indicated at right. Molecular weight ($M_W$) markers are indicated at left. Results are representative of at least three biological replicates. (c) Glycosylation efficiency of cell-based scFv-HER2 glycosylation where % glycosylated was calculated as the ratio g1/[g0+g1], and g0 and g1 values were determined by densitometric quantification of bands in anti-His immunoblots in (b) for each neoglycoprotein.

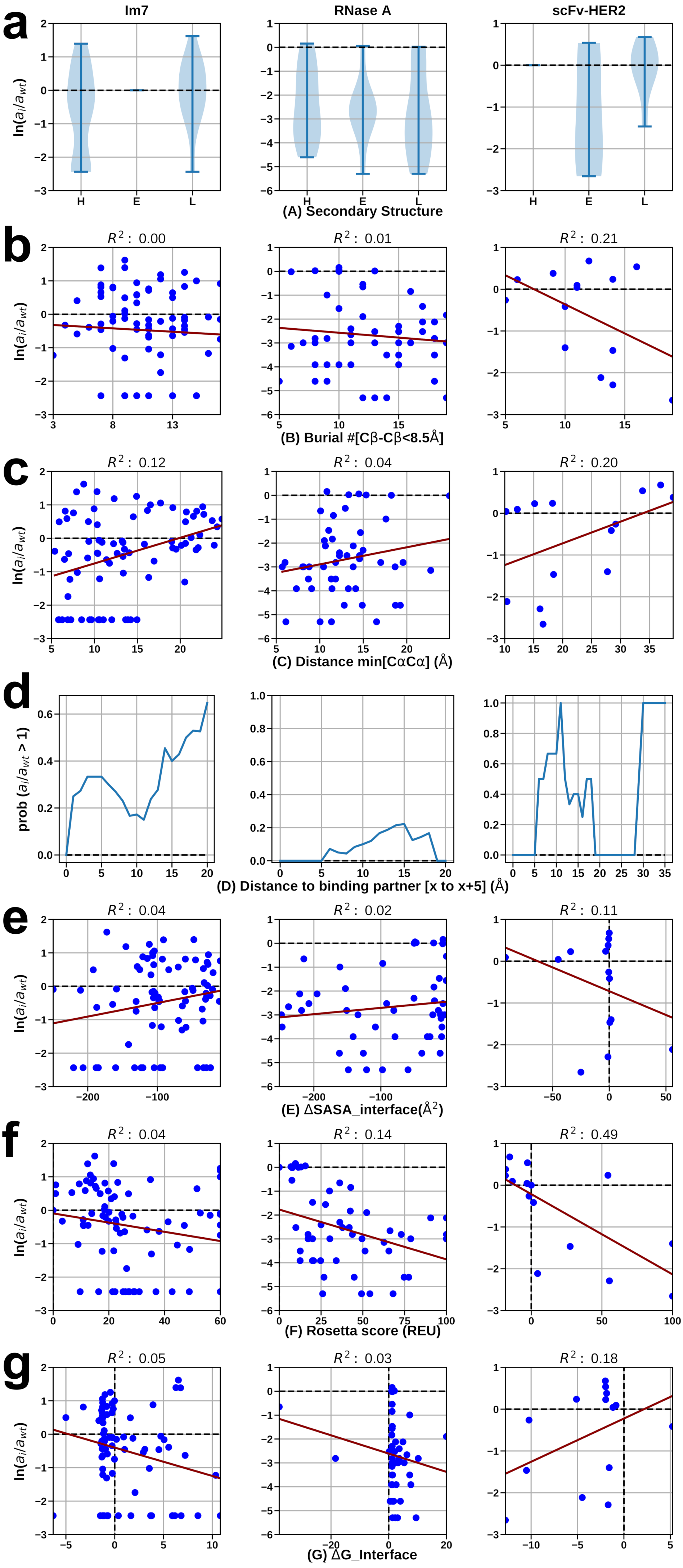


**Supplementary Figure 9. Computational analyses for aglycosylated variants.** Geometric measures and Rosetta calculated energies plotted against activity of aglycosylated Im7, RNase A, and scFv-HER2 variants as indicated. For Im7 and scFv-HER2, activity represents binding activity for aglycosylated variants; for RNase A, activity represents catalytic activity for aglycosylated variants. Activity was normalized by activity of wild-type (wt) protein. For variants where no activity was detected, activity was plotted using the instrument detection limit.

(a) Distribution of natural log of the ratio of activities for sequon substituted variants relative to wt in different secondary structure elements (H, E, and L denote α-helix, β-strand and loop, respectively).

(b) Burial of sequon (measured by number of Cβ residues within 8.5 Å of the central sequon residue Cβ) versus natural log of the activity ratio.

(c) Minimum distance of sequon center residue to any residue of binding partner (for RNase A, the distance between His119 and the center residue of substituted sequon) versus the natural log of the activity ratio.

(d) Distribution of probability of increased activity in the variant relative to wt given the distance of the sequon position to the binding partner (or His119 for RNase A).

(e) Change in solvent accessible surface area buried upon binding ColE7, RNA, or HER2-ED (relative to wt) versus the natural log of the activity ratio.

(f) Change in Rosetta total score of variants (relative to wt) versus the natural log of the activity ratio. Rosetta scores over 100 REU are shown at 100 REU.

(g) Change in Rosetta binding energy (relative to wt) versus the natural log of the activity ratio.

In all graphs, red lines indicate regressions, with the $R^2$ values listed above each plot.

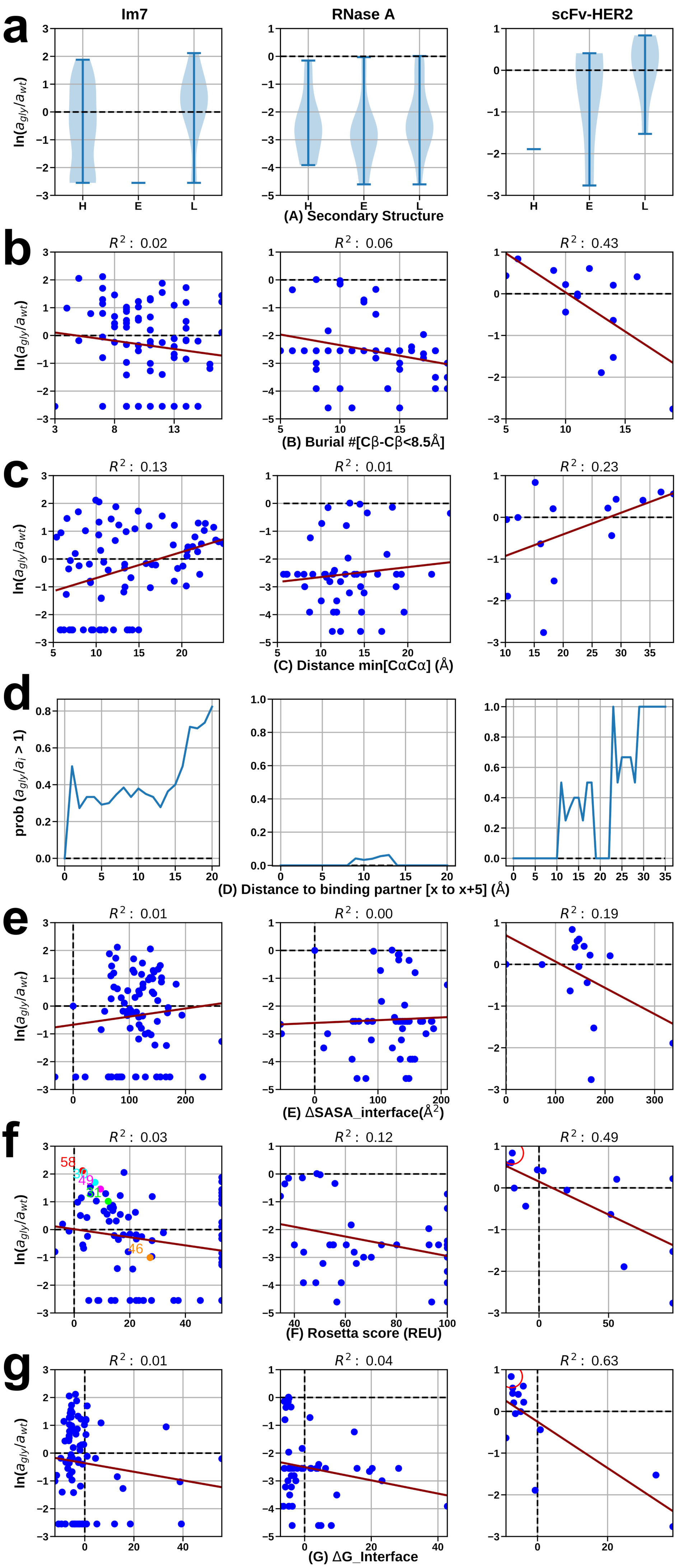


**Supplementary Figure 10. Computational analyses for glycosylated variants.** Geometric measures and Rosetta calculated energies plotted against activity of glycosylated Im7, RNase A, and scFv-HER2 variants as indicated. For Im7 and scFv-HER2, activity represents binding activity for glycosylated variants; for RNase A, activity represents catalytic activity for glycosylated variants. Activity was normalized by activity of wild-type (wt) protein. For variants where no activity was detected, activity was plotted using the instrument detection limit.

(a) Distribution of natural log of the ratio of activities for sequon substituted variants relative to wt in different secondary structure elements (H, E, and L denote α-helix, β-strand and loop, respectively).

(b) Burial of sequon (measured by number of Cβ residues within 8.5 Å of the central sequon residue Cβ) versus natural log of the activity ratio.

(c) Minimum distance of sequon center residue to any residue of binding partner (for RNase A, the distance between His119 and the center residue of substituted sequon) versus the natural log of the activity ratio.

(d) Distribution of probability of increased activity in the variant relative to wt given the distance of the sequon position to the binding partner (or His119 for RNase A).

(e) Change in solvent accessible surface area buried upon binding ColE7, RNA, or HER2-ED (relative to wt) versus the natural log of the activity ratio.

(f) Change in Rosetta total score of variants (relative to wt) versus the natural log of the activity ratio. Rosetta scores over 100 REU are shown at 100 REU.

(g) Change in Rosetta binding energy (relative to wt) versus the natural log of the activity ratio.

In all graphs, red lines indicate regressions, with the $R^2$ values listed above each plot.

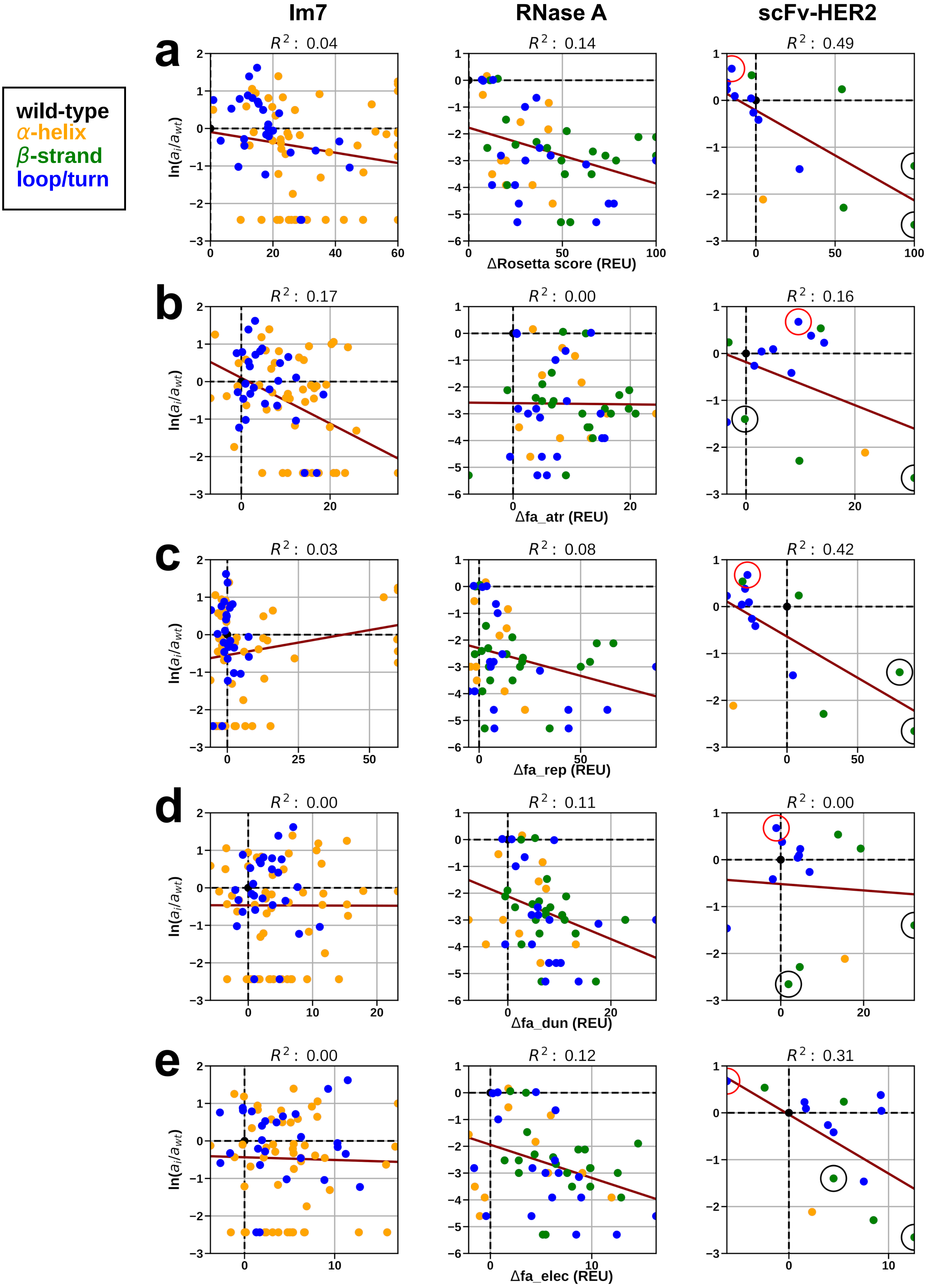


**Supplementary Figure 11. Rosetta score decomposition for aglycosylated variants.** Decomposed Rosetta score terms plotted against aglycosylated activity for Im7, RNase A and scFv-HER2 variants as indicated. For Im7 and scFv-HER2, activity represents binding activity for aglycosylated variants; for RNase A, activity represents catalytic activity for aglycosylated variants. Activities were normalized to the wild-type (wt) protein. (a) Change in Rosetta total score of aglycosylated variants (relative to wt) plotted versus log activity ratio. (b) Change in attractive Lennard-Jones score (fa_atr in Rosetta) versus log activity ratio. (c) Change in repulsive Lennard-Jones score (fa_rep) versus log activity ratio. (d) Change in Dunbrack rotamer score (fa_dun) versus log activity ratio. (e) Change in Coulombic electrostatic potential versus log activity ratio. Circles in scFv-HER2 column identify residues discussed in main text: N36 $V_L$, N113 $V_H$: black; N108 $V_L$: red.

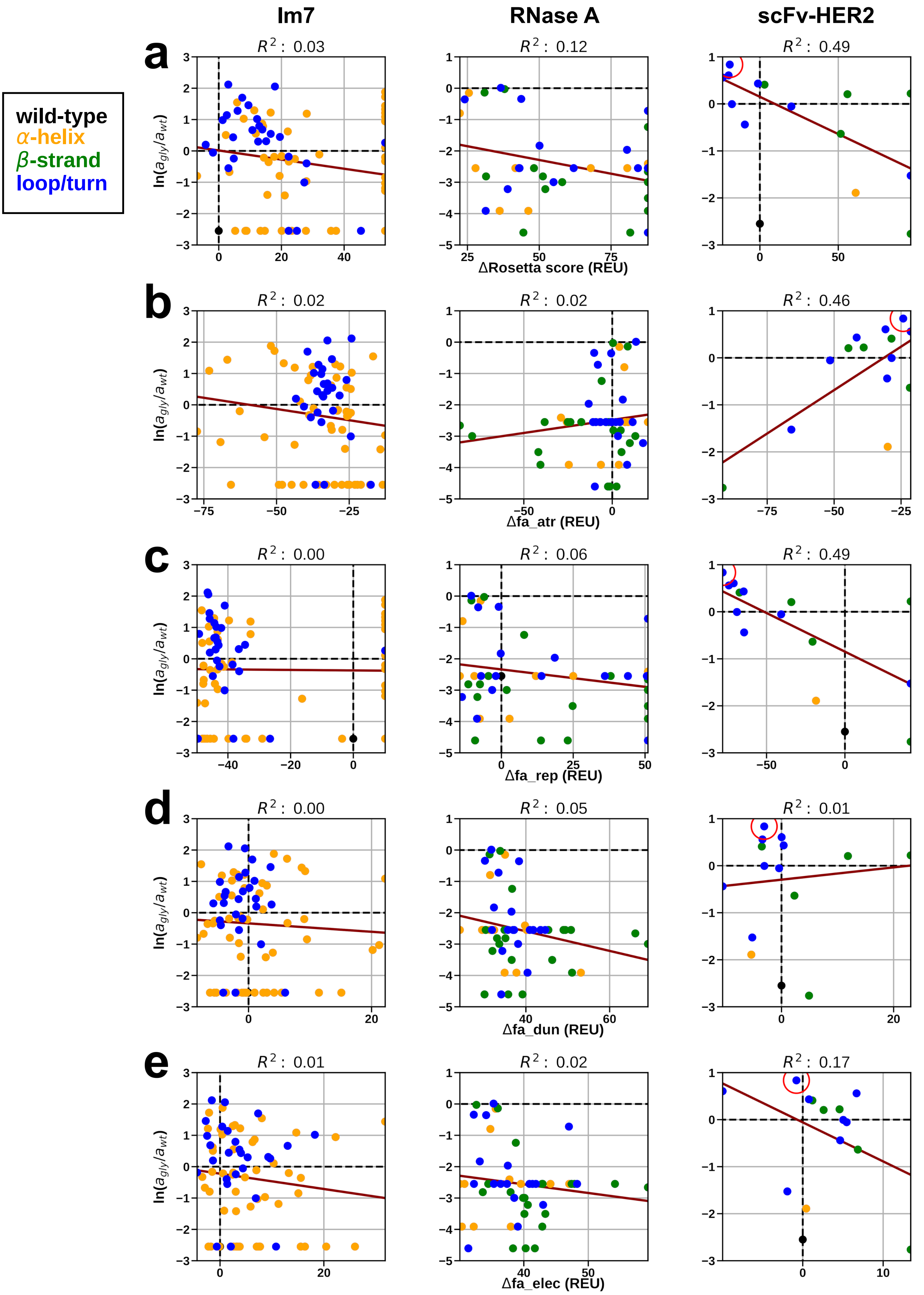


**Supplementary Figure 12. Rosetta score decomposition for glycosylated variants.** Decomposed Rosetta score terms plotted against glycosylated activity for Im7, RNase A and scFv-HER2 variants as indicated. For Im7 and scFv-HER2, activity represents binding activity for glycosylated variants; for RNase A, activity represents catalytic activity for glycosylated variants. Activities were normalized to the wild-type (wt) protein. (a) Change in Rosetta total score of glycosylated variants (relative to wt) plotted versus log activity ratio. (b) Change in attractive Lennard-Jones score (fa_atr in Rosetta) versus log activity ratio. (c) Change in repulsive Lennard-Jones score (fa_rep) versus log activity ratio. (d) Change in Dunbrack rotamer score (fa_dun) versus log activity ratio. (e) Change in Coulombic electrostatic potential versus log activity ratio. Red circle in scFv-HER2 column identifies residue N58 $V_L$ discussed in main text.

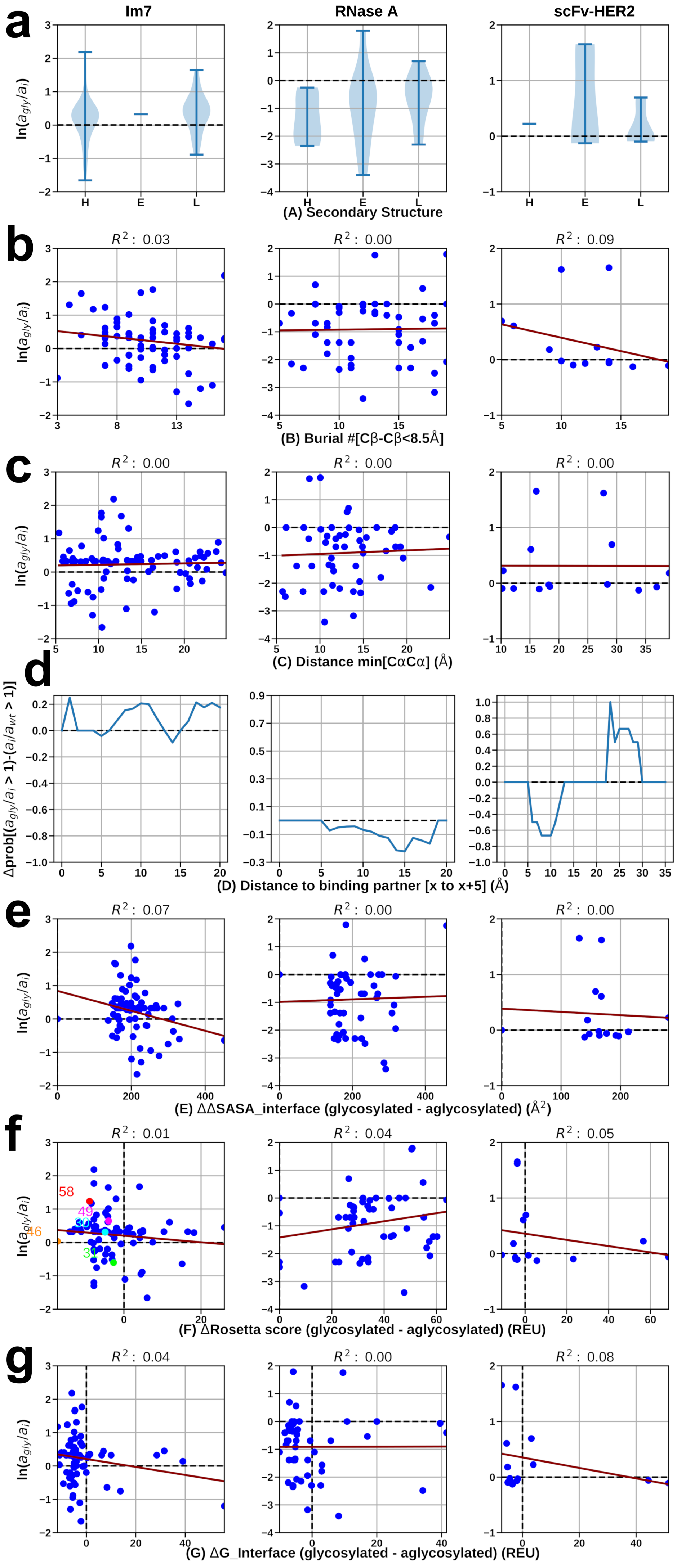


**Supplementary Figure 13. Computational analyses comparing glycosylated and aglycosylated variants.** Geometric measures and Rosetta calculated energies plotted against the glycosylated/aglycosylated activity ratio for Im7, RNase A, and scFv-HER2 variants as indicated. For Im7 and scFv-HER2, activity ratio represents binding activity for glycosylated/aglycosylated variants; for RNase A, activity ratio represents catalytic activity for glycosylated/aglycosylated variants.

(a) Distribution of natural log of the activity ratio of glycosylated and aglycosylated variants in secondary structure elements (H, E, and L stand for α-helix, β-strand and loop, respectively).

(b) Burial of sequon (measured by number of Cβ residues within 8.5 Å of the central sequon residue Cβ) versus natural log ratio of activities.

(c) Minimum distance of sequon center residue to any residue of binding partner (for RNase A, the distance between His119 and the center residue of substituted sequon) versus the log ratio of activity.

(d) Distribution of probability of increased activity in the glycosylated variant relative to the aglycosylated variant given the distance of the sequon position to the binding partner (or His119 for RNase A).

(e) Difference in solvent accessible surface area between glycosylated and aglycosylated variants buried upon binding ColE7, RNA, or HER2-ED versus natural log of the activity ratio.

(f) Difference in Rosetta total score from glycosylated to aglycosylated variants versus their respective natural log of activity ratio. Rosetta scores over 100 REU are shown at 100 REU.

(g) Difference in Rosetta binding energy between glycosylated and aglycosylated variants versus respective natural log of activity ratio.

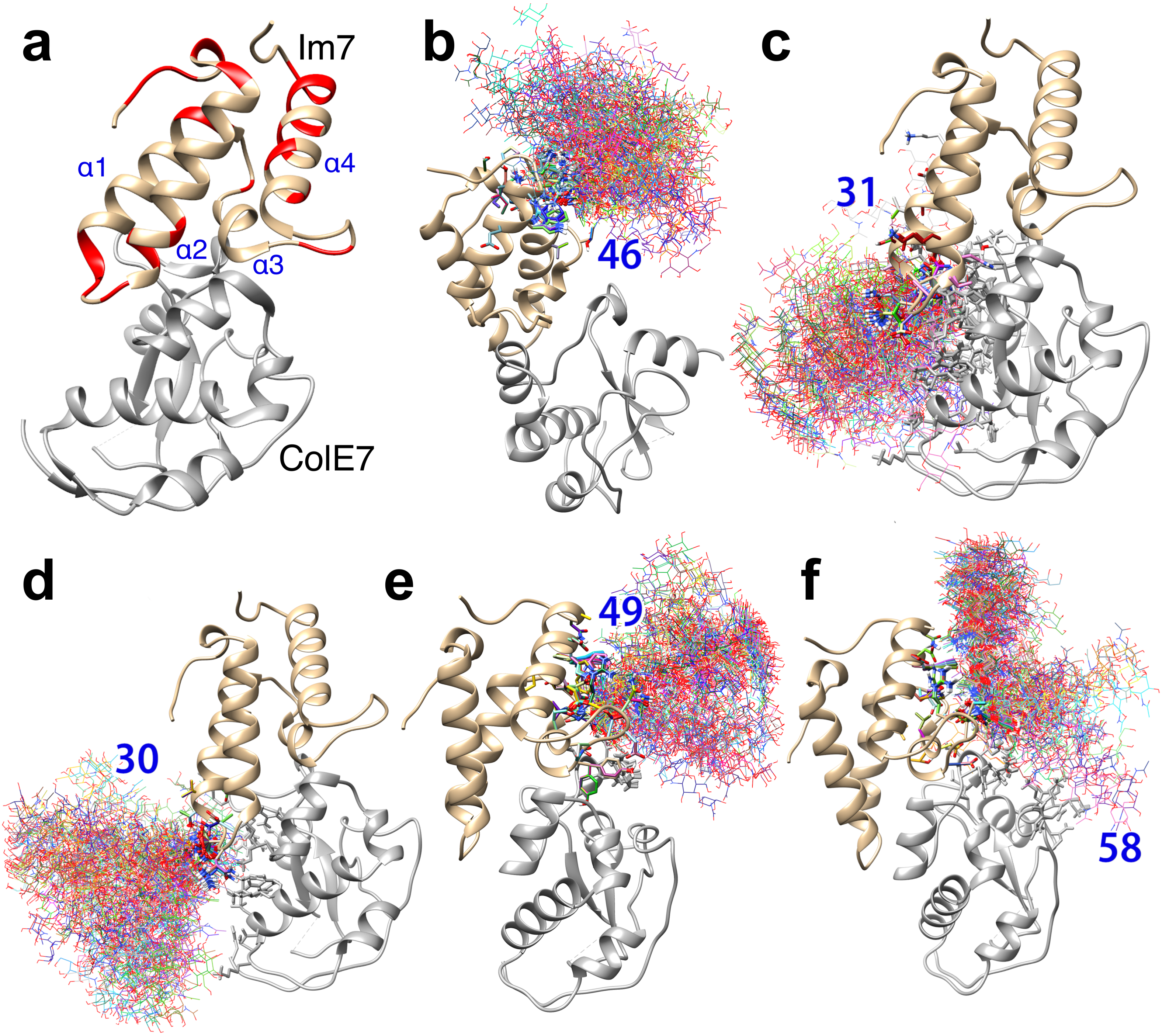


**Supplementary Figure 14. Effect of sequon substitution and glycosylation on Im7-ColE7 binding.** Im7 (gold) in complex with ColE7 (gray). (a) Sites on Im7 with stronger binding than wt after sequon substitution (red). (b-f) Ensemble of glycosylated mutants 46, 31, 30, 49 and 58 ($a_{gly}$ / $a_i$ = 1.04, 0.55, 1.36, 1.91, and 3.44, respectively) showing low-energy conformations of conjugated glycans. *N*-glycans are shown as lines with oxygens in red, nitrogens in blue, and carbons in different colors for each model; Im7 and ColE7 side chains that interact with the *N*-glycan are shown as sticks.
